# Transcriptomic analysis of meiotic genes during the mitosis-to-meiosis transition in *Drosophila* females

**DOI:** 10.1101/2024.07.10.602987

**Authors:** Ana Maria Vallés, Thomas Rubin, Nicolas Macaisne, Laurine Dal Toe, Anahi Molla-Herman, Christophe Antoniewski, Jean-René Huynh

## Abstract

Germline cells produce gametes, which are specialized cells essential for sexual reproduction. Germline cells first amplify through several rounds of mitosis before switching to the meiotic program, which requires specific sets of proteins for DNA recombination, chromosome pairing and segregation. Surprisingly, we previously found that some proteins of the synaptonemal complex, a prophase I meiotic structure, are already expressed and required in the mitotic region of *Drosophila* females. Here, to assess if additional meiotic genes were expressed earlier than expected, we isolated mitotic and meiotic cell populations to compare their RNA content. Our transcriptomic analysis reveals that all known meiosis I genes are already expressed in the mitotic region, however, only some of them are translated. As a case study, we focused on *mei-W68*, the *Drosophila* homologue of *Spo11*, to assess its expression at both the mRNA and protein levels, and used different mutant alleles to assay for a pre-meiotic function. We could not detect any functional role for Mei-W68 during homologous chromosome pairing in dividing germ cells. Our study paves the way for further functional analysis of meiotic genes expressed in the mitotic region.

**Article Summary:** Germline cells, crucial for sexual reproduction, were thought to switch to meiosis only after several rounds of mitosis. Surprisingly, a few meiotic proteins were found active in the mitotic phase of female flies. Here, we discovered that all known meiosis genes were expressed during mitosis, but only some produced proteins. This study suggests that genes related to reproduction are active earlier than expected, prompting further exploration into their functions during early cell division.

## Introduction

In organisms reproducing sexually, germline cells produce oocytes and sperms as gametes. Germline cell differentiation starts by an amplification phase through mitosis to increase their numbers and create a pool of precursor cells (Cinalli *et al*. 2008). They then switch to meiosis, which comprises two rounds of nuclear divisions to produce haploid gametes. Meiosis is specific to germline cells and requires the expression of unique molecular machineries to pair, recombine and segregate homologous chromosomes. How germline cells switch from a mitotic to a meiotic program is not fully understood.

Meiosis starts by an extended prophase I during which homologous chromosomes have to find each other in the nuclear space to pair (Bhalla and Dernburg 2008; Zickler and Kleckner 2015). Once homologous chromosomes are paired, their association is stabilized by the synaptonemal complex (SC), the proteinaceous structure that holds homologous axes together (synapsis) and promotes genetic recombination (Cahoon and Hawley 2016). Recombination starts by the formation of developmentally programmed double-strand breaks (DSBs), which can be later repaired as crossovers. Meiotic DSBs are induced by the topoisomerase-like Spo11, which is conserved in all species (Keeney *et al*. 1997; De massy 2013). These chromosome exchanges create physical links called chiasmata, which keep homologues paired until they orient towards opposite poles of the spindle. This period is subdivided in five classical stages (leptotene, zygotene, pachytene, diplotene, and diakinesis) based on chromosome morphologies. The initiation of the pairing process has been defined at the early zygotene stage in *Saccharomyces cerevisiae* (Tsubouchi T 2005) and at the leptotene stage in *C. elegans* (CRITTENDEN SL 2006; Rohozkova *et al*. 2019) zebrafish (Blokhina *et al*. 2019) and mice (Ishiguro *et al*. 2014; Scherthan *et al*. 2014), by FISH analysis and chromosome axis protein imaging. Moreover, chromosome movements, forces and molecular players that promote pairing have been well characterized by live imaging microscopy in these species (Rubin *et al*. 2020; Kim *et al*. 2022).

However, we and others have found that homologous chromosomes start to pair through centromeres and euchromatic loci during the mitotic phases preceding leptotene in both *Drosophila* males and females (Cahoon and Hawley 2013; CHRISTOPHOROU N 2013; JOYCE EF 2013; Christophorou *et al*. 2015; Rubin *et al*. 2022). Moreover, we showed that this pre-meiotic pairing requires components of the synaptonemal complex, a structure specific to prophase I of meiosis (CHRISTOPHOROU N 2013; Rubin *et al*. 2022). Indeed, the C(3)G and Corona proteins, which form the central region of the SC, are transcribed and translated in the mitotic region and localize on one side of centromeres. It is similar to the initiation of meiosis in budding yeast, where centromeres become “coupled” before meiotic prophase (Tsubouchi T 2005). This early association also depends on Zip1, a central component of the SC functionally similar to C(3)G in flies and SYCP1 in mice. Furthermore, recent analyses in mice have shown that meiotic genes involved in prophase I are expressed and translated long before the initiation of the meiotic process (Wang *et al*. 2001; Evans *et al*. 2014; Zheng *et al*. 2022). For example, the meiotic cohesin REC8, as in *C. elegans* (Pasierbek *et al*. 2001) and synaptonemal complex proteins are expressed and actively translated in spermatogonia, which go through several mitotic divisions before meiotic entry. In addition, Spo11 protein is also found at very low levels in spermatogonia (Fang *et al*. 2021).

Here, to assess if additional meiotic genes were expressed in the mitotic region of *Drosophila* females, we analyzed the whole genome transcriptome of mitotic and meiotic germline cells.

## Results

### 1. Meiotic genes are expressed in the mitotic region

In *Drosophila* females, the processes of mitosis and meiosis occur sequentially throughout the adult life in a structure called the germarium located at the tip of each ovary (SPRADLING 1993). At the anterior-most part is the mitotic zone, also known as region 1. In this zone, germline stem cells (GSCs) proliferate and self-renew by receiving signals from adjacent somatic tissue that induce the expression of stem cell promoting factors like *nanos*, which mediate the translational repression of differentiation genes (Slaidina and Lehmann 2014). GSCs divide mostly asymmetrically and generate a posterior daughter cell, which differentiates into a precursor cell called cystoblast (CB). The CB undergoes four rounds of mitosis, resulting in the formation of a germline cyst consisting of 16 cells (Figure 1A) (Huynh and St Johnston 2004). During these mitotic divisions, cells remain connected through ring canals and a specialized organelle called the fusome. The branching pattern of the fusome is a useful marker for distinguishing the different stages within the mitotic zone, i.e. GSCs, CBs, and cysts of 2, 4, 8, and 16 cells (de Cuevas and Spradling 1998). The period of rapid synchronized divisions marks the transition phase and the commitment to differentiation. The Bag of marble (Bam) protein induces the differentiation of CBs and its expression is spatially restricted: suppressed by self-renewal factors in GSCs and activated in CBs and 2, 4, 8-cell cysts (Fig. 1B and C) (Chen and Mckearin 2003). After the last mitosis, cysts enter the meiotic zone, also known as region 2a, where all 16 cells that look identical enter meiosis (Carpenter 1975). The presence of the synaptonemal complex (SC) in this early meiotic zone marks the initiation of prophase I, with only two pro-oocytes progressing to form a complete SC (Hughes *et al*. 2018). At this stage, meiotic double-strand breaks (DSBs) are induced. As the cyst reaches region 2b, only one cell within the cyst will become an oocyte, while the remaining 15 cells develop into nurse cells and undergo DNA endoreplication. In this region, the cyst undergoes a significant morphological change, adopting a disc-like shape that is one cell thick and spans the entire width of the germarium. Concomitantly, somatic follicle cells begin to migrate and enclose the cyst. As the cyst advances to region 3, also known as stage 1, it assumes a rounded shape forming a sphere. At late pachytene, the oocyte stage is marked with SC and consistently positioned at the posterior pole. Subsequently, the cyst exits the germarium and enters the vitellarium (Huynh and St Johnston 2004).

**Figure 1.**
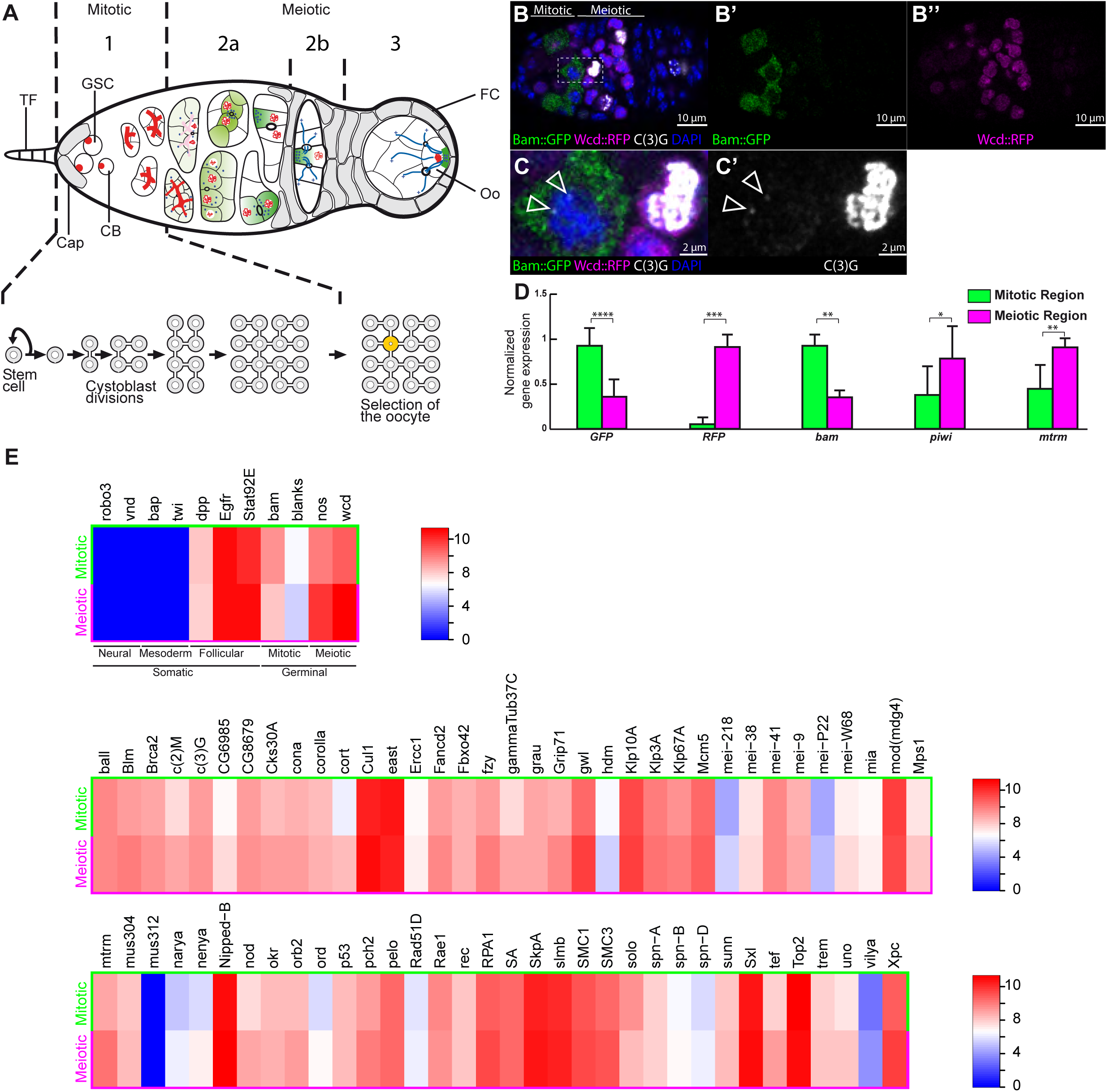
Meiotic genes are expressed in the mitotic region of the *Drosophila* germarium. (A) Drosophila germarium depicting the mitotic and meiotic regions. In the anterior part (Mitotic zone, also called region 1) at the base of the terminal filament (TF), somatic cap cells surround germline stem cells (GSC) that divide four times giving rise to a 16-cells cyst. GSCs and cystoblasts (CB) are marked by the spectrosome (red circle), and the developing two-, four, and eight-cell cysts by the fusome (red-branched structure). After the last mitosis, cysts move to the meiotic zone, subdivided in region 2a, 2b and 3. Early in region 2a, the synaptonemal complex (red thin lines) marks the pro-oocytes with four ring canals. By region 2b, the oocyte is selected and is the only cell (yellow) to remain in meiosis. The follicle cells (FC) start to migrate and surround the germline cells as the cyst moves posteriorly to region 3. (B-B’’) Confocal Z-section of a germarium labelling the mitotic region with Bam::GFP (Green), the meiotic region with Wcd::RFP (magenta), the synaptonemal complex with C(3)G (white) and cell nuclei with DAPI (blue). (C, C’) Magnification of hatched square in B showing C(3)G nuclear labelling of a cell in the mitotic region (open arrows) adjacent to a synaptonemal complex labelled pro-oocyte. Scale bar: 10 μm in B-B’’ 2 μm in C, C’. (D) RT-PCR gene expression levels of FACS-separated mitotic (green) and meiotic (magenta) cells using primers to GFP, RFP, *bam*, *piwi* and *mtrm*. Gene expression levels are defined to 1 relative to the highest value within both population after rp49 normalization. * p ≤ 0.05, ** p ≤ 0.01, *** p ≤ 0.001, **** p ≤ 0.0001 (Mann-Whitney U-test) (E) Heat map of known meiotic genes expressed in FACS-separated Bam::GFP (Mitotic) and Wcd::RFP cells (Meiotic). In the upper pannel are the somatic genes, *robo3, vnd* (neural); *bap, twi (*mesodermal); *dpp, Egfr, Stat92e* (follicle cells) and the sorting controls *bam, blanks, nos* and *wcd*. The middle and lower panels represent the heat map of meiotic genes. Scale represents log2 expression gradient for genes expressed in each of the two regions. Notice that neural and mesodermal contaminants are not detected while follicle cells ones are equally present in both cell populations.

Although meiosis is described as beginning in early region 2a of the germarium, several proteins needed for homologous chromosomes pairing are already present in mitotically dividing cells of region 1. The synaptonemal complex protein C(3)G is one example, which localizes near the centromeres of chromosomes II and III and whose expression is required for initiating centromeric pairing ((CHRISTOPHOROU N 2013); Figure 1 B,C). To gain a more exhaustive view of the spatiotemporal expression of meiotic genes, we separated the mitotic and meiotic cell populations by FACS and then processed the RNA for high-throughput sequencing. The separation method relied on the restrictive expression pattern and properties of Bam and Wcd transgenic proteins (Valles and Huynh 2020): Bam::GFP is detectable only in 2-8 cell cysts of region 1 and was used to label the mitotic region (Mckearin and Ohlstein 1995). Wcd::RFP has a fast turnover and, when driven by *nanos*-Gal4, it labels a few GSCs, and mostly region 2a/b cells; we therefore used it to identify cells in the first stages of meiosis I. The transgenic line (Bam::GFP; nos>Wcd::RFP) labeled germaria and allowed efficient separation of both mitotic and meiotic germ cell population (see Material and Methods section Figure S1-S3; Table S5). We confirmed the efficiency of cell-sorting by qRT-PCR for specific transcripts. We found that GFP cells were strongly enriched in GFP and *bam* RNA transcripts, while RFP cells were enriched in RFP and *wcd* RNAs (Figure 1D, E). Endogenous Piwi protein was shown to be strongly downregulated in 2- to 8-cell cysts forming a “piwilesspocket (pilp)” (Dufourt *et al*. 2014). Similarly, we found that *piwi* mRNA levels were lower in the mitotic region compared to the meiotic region (Figure 1D). *matrimony* (*mtrm*) was reported to be very lowly expressed in the mitotic region and higher in the meiotic region by different methods such as single-cell RNAseq and synchronized germline cells (Slaidina *et al*. 2021; Samuels *et al*. 2024). We confirmed these results by RT-qPCR and RNAseq (Figure 1D and 1E; Table S1-4). As an additional control, we used *blanks*, as this gene was previously shown to display the opposite trend with higher expression in mitotic cells than in meiotic cells (Slaidina *et al*. 2021; Samuels *et al*. 2024). Similarly, our results indicate that *blanks* expression levels are higher in the GFP+ cell population than in the RFP+ population (Figure 1E). To evaluate the contamination by other cell types, we searched for somatic cell markers such as *Robo3*, *vnd* (neural cell), *twist* and *bap* (mesoderm) and *dpp*, *egfr*, *STAT92*, which are expressed in somatic cells in the germarium but not germline cells. We found that *Robo3, vnd, twist* and *bap* RNAs were absent from both cell populations, however, we found that *dpp*, *egfr* and *Stat92E* were equally present in GFP+ and RFP+ cells (Figure 1E and Table S1-2). These results indicated that there was no contamination by neural or mesodermal tissues, but that some ovarian somatic cells were equally present in both isolated cell populations. Despite the presence of somatic cells in both samples, our control experiments demonstrated that we were able to separate germline mitotic cells from meiotic cells, and that our results were consistent with previously published data.

We then took advantage of these transcriptome datasets to focus on genes required for the initial stages of meiosis I. We used the single GO term “meiosis I” in Flybase and removed male-specific genes to identify 69 genes. We found that all of these genes were expressed in both mitotic and meiotic cell populations (Figure 1E, Table S1-2). As previously shown by antibody staining, the synaptonemal complex components C(3)G, Corona and Ord were all found expressed in the mitotic region. We used the DESeq2 package to analyze the differential expression between these genes in the mitotic and meiotic cell populations. Except for a few exceptions, we found that most meiotic genes were expressed at low levels in region 1, and that their expression increase on average by 1.56 folds in region 2 (Figure 1E, Table S1). At one end, *hdm* expression is downregulated 2.3 folds from mitosis to meiosis, almost as strongly as our control gene *bam*, which expression is decreased by 2.7 folds (Figure 1E, Table S1). At the other end, the expression of *cortex* is increased by 4.1 folds. Genes encoding proteins required for homologous recombination such as *mei-W68* and *mei-P22* were among the least expressed in both cell types; nevertheless, their expression increased by 1.5 and 1.6 folds respectively in region 2.

To further validate our results, we performed a highly sensitive in situ hybridization (FISH) using the Hybridization Chain Reaction (HCR) method for *C(3)G*, *Nipped-B* and *mei-W68* RNAs (Choi *et al*. 2018). To unambiguously distinguish the different stages within the mitotic and meiotic regions (Figure 2, yellow dotted line), we labeled the fusome with an antibody against α-Spectrin (Figure S2A-B). Consistent with our RNAseq data, we found that all three genes were expressed in region 1 cells; at very low levels for *mei-W68* and higher levels for *C(3)G* and *Nipped-B* (Figure 2A, C, E). Quantification of the fluorescent signals also revealed an increase in RNA levels for all three genes as found by the RNAseq analysis (Figure 2B, D, F).

**Figure 2.**
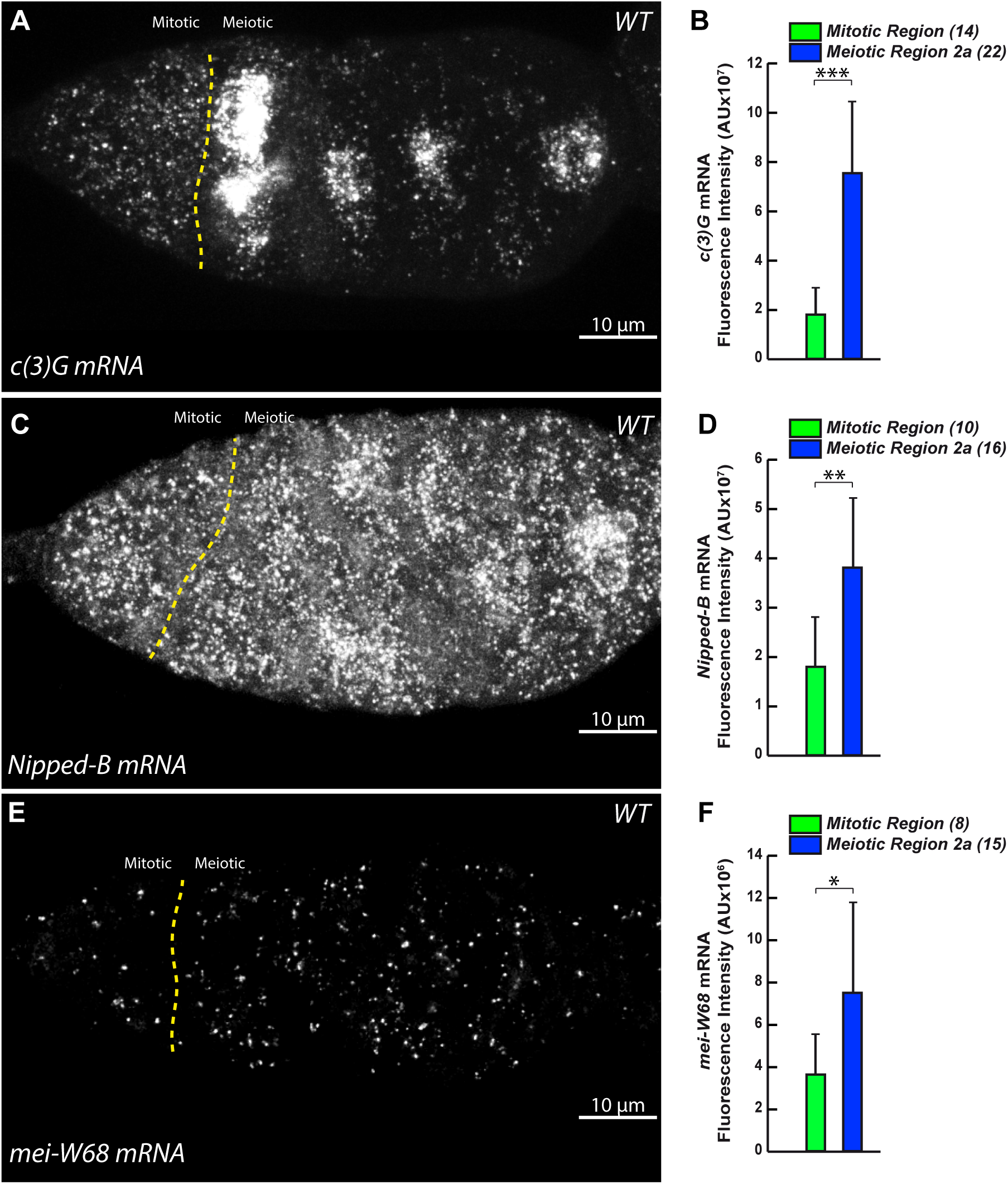
*c(3)G*, *Nipped-B* and *mei-W68* meiotic genes mRNA are detected in the mitotic region and their levels increase in the meiotic region. (A) Confocal Z-section projection of a *WT* germarium labelled for *c(3)G* mRNA by HCR *in situ* hybridization. The yellow dashed line delimits the boundary of mitotic and meiotic regions. Scale bar: 10 μm. (B) Graph plots *c(3)G mRNA* Fluorescence Intensity in mitotic and meiotic 2a regions. *** p ≤ 0.001 (Mann-Whitney U-test). (C) Confocal Z-section projection of a *WT* germarium labelled for *Nipped-B* mRNA by HCR *in situ* hybridization. The yellow dashed line delimits the boundary of mitotic and meiotic regions. Scale bar: 10 μm. (D) Graph plots *Nipped-B mRNA* Fluorescence Intensity in mitotic and meiotic 2a regions. ** p ≤ 0.01 (Mann-Whitney U-test). (E) Confocal Z-section projection of a *WT* germarium labelled for *mei-W68* mRNA by HCR *in situ* hybridization. The yellow dashed line delimits the boundary of mitotic and meiotic 2a regions. Scale bar: 10 μm. (F) Graph plots *mei-W68 mRNA* Fluorescence Intensity in mitotic and meiotic regions. * p ≤ 0.05 (Mann-Whitney U-test). (*n*) is the number of germaria analyzed for each probe.

Overall, we concluded that we were able to isolate the mitotic germline cells from the meiotic cells, and that all meiotic genes started to be expressed in mitotic cells.

### 2. *mei-W68* gene is expressed in the mitotic region but Mei-W68 protein is only detected in meiotic cells

Spo11 and TopoVIBL form a meiosis-specific complex, which is conserved across species. In *Drosophila*, Mei-W68 is the homologue of Spo11, while Mei-P22 is a potential homologue of TopoVIBL (Robert *et al*. 2016; Vrielynck *et al*. 2016). The conserved function of this complex is to generate DSBs to initiate recombination between homologous chromosomes. However, in some species such as mouse, zebrafish, and recently jellyfish, these DSBs are also required for the formation of a SC (Romanienko and Camerini-Otero 2000; Blokhina *et al*. 2019; Munro *et al*. 2023), whereas it is not the case in *C. elegans* and *Drosophila* females (Dernburg *et al*. 1998; Mckim *et al*. 1998). Here, we wanted to test whether Mei-W68 played a role in homologous chromosome pairing in region 1 before the initiation of meiotic DSBs.

As shown above using RNAseq and RNA FISH, we found that *mei-W68* mRNA is present at low levels in region 1. Next, we wanted to examine whether Mei-W68 protein was present in region 1. Since there is no antibody against Mei-W68 available in *Drosophila* and that Spo11 homologues are also very hard to detect in other species, we decided to knock-in a small 3xHA-6xHis tag by CRISPR-Cas9 at the C-terminus of the endogenous protein (Figure S4C). Despite successful integration, we found that the fusion protein was not functional, as no DSBs could be detected with an anti-γH2Av antibody in mei-W68-HA flies (Figure S5). Furthermore, we found that the frequencies of X and chromosome II non-disjunction were similar in *mei-W68-HA*/*Df(*BSC782) and *mei-W68^1^/Df(*BSC782) (Table S6A, S6B). However, *mei-W68-HA* RNAs were nonetheless translated as we were able to detect a specific signal in region 2a using an anti-HA antibody (Figure 3B). Quantification of this signal revealed that the levels of Mei-W68-HA protein in region 1 were at background levels, and dramatically increased in meiotic cells (Figure 3A’, B’, C). These results showed that Mei-W68 protein is probably not present in region 1, and that *mei-W68* mRNA is translated only in *Drosophila* meiotic cells.

**Figure 3.**
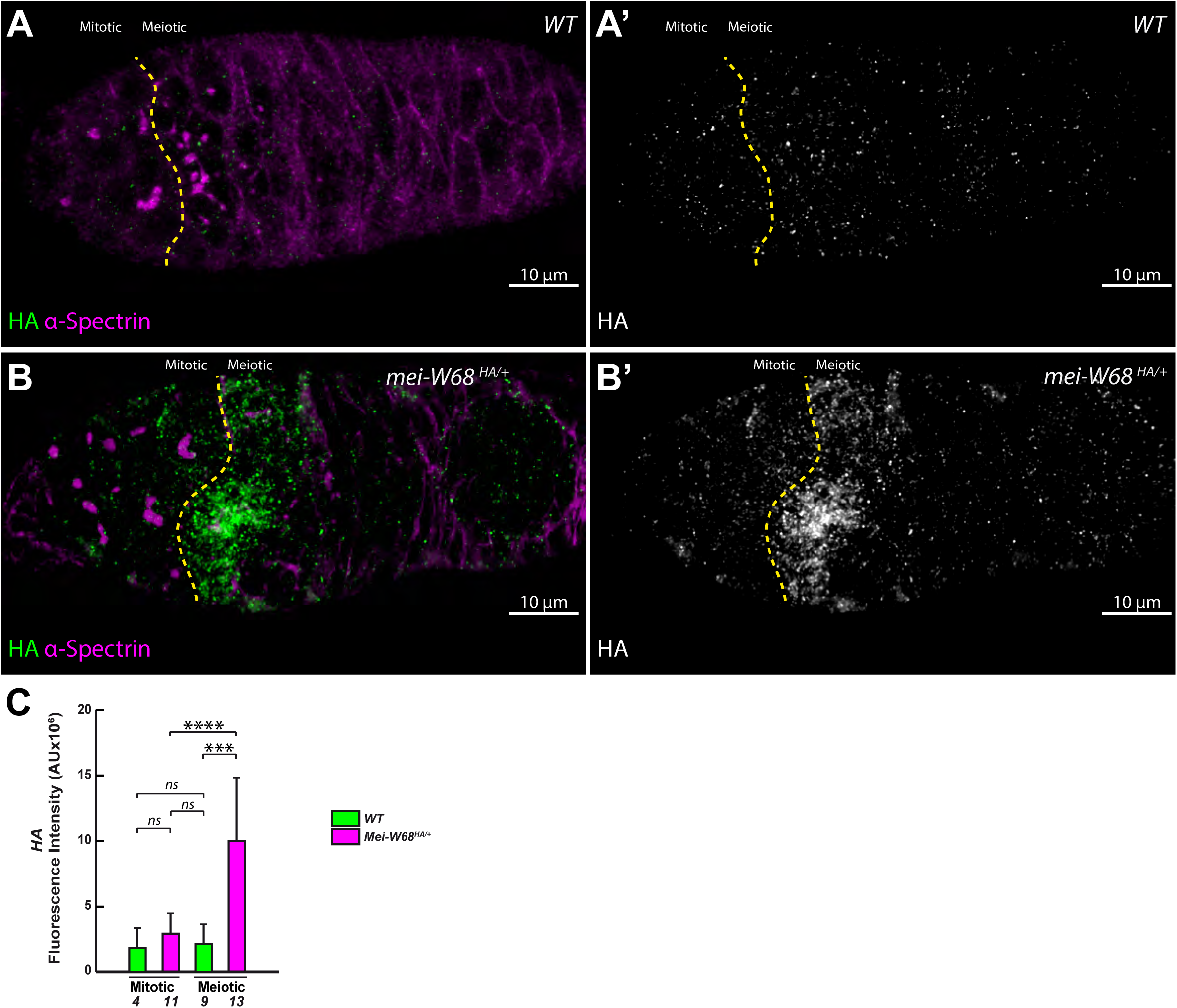
Mei-W68 protein is only detected in the meiotic region. (A-A’) Confocal Z-section projection of a *WT* germarium immunostained for HA (green) and the fusome (magenta). The yellow dashed line delimits the boundary of mitotic and meiotic regions. Scale bar: 10 μm. (B, B’) Confocal Z-section projection of a *mei*-*W68^HA^/+* germarium immunostained for HA (green) and the fusome (magenta). The yellow dashed line delimits the boundary of mitotic and meiotic regions. Scale bar: 10 μm. Note that HA immunostaining is barely detectable in both regions (A, A’) of *WT*, while HA is clearly confined to the meiotic region of *W68^HA^/+* (compare mitotic and meiotic region in in B, B’). (C) Graph plots HA Fluorescence Intensity in mitotic and meiotic regions. ns p ≥ 0.05, *** p ≤ 0.001, **** p ≤ 0.0001 (Mann-Whitney U-test). Numbers below bars represent the germaria analysed.

### 3. Mei-W68 and Mei-P22 are dispensable for centromere pairing in the mitotic region

The failure to detect Mei-W68 protein in the mitotic region could be due to limitations in our detection methods combined with its low expression levels, as suggested by our transcriptome analysis. We therefore used a functional assay to test for a requirement of Mei-W68 in region 1. In mouse germline cells, Spo11 has been proposed to be required for pre-meiotic pairing of homologous chromosomes (Boateng *et al*. 2013). We thus assayed whether Mei-W68 was required for pre-meiotic pairing of centromeres in region 1. To this aim, we used three different mutant conditions. Firstly, in *mei-W68^1^/DfBSC782* mutant germaria, there is a 5kb insertion of a transposable element in the first exon, and there is likely no protein made (Mckim and Hayashi-Hagihara 1998) (Figure S4A). Secondly, we replaced by CRISPR-Cas9 the endogenous locus with a form of *mei-W68* mutated in the catalytic domain. Based on sequence alignment of similar constructs in yeast and mouse, we replaced two Tyrosine (Y80 and Y81) by two Phenylalanine in the catalytic domain (*mei-W68^CD^* Figure S4B) (Diaz *et al*. 2002; Boateng *et al*. 2013). In this mutant background (*mei-W68^1^/DfBSC782* and *mei-W68^CD^*) no DSBs could be detected with an anti-γH2Av antibody (Figure S6A, S6B, S6C). Furthermore, we found high levels of non-disjunction for both the X and second chromosomes (Table S6A, S6B), indicating that *mei-W68^CD^* is a strong mutant allele of *mei-W68.* Thirdly, we used a *mei-P22^P22^* mutant allele and confirmed that DSBs were also completely absent (Figure S6D).

In previous studies, we observed that centromere pairing became prominent in 8-cell germline cysts (CHRISTOPHOROU N 2013). *Drosophila* diploid cells have four pairs of homologous chromosomes, resulting in eight chromosomes per cell. When all homologues are paired, we can observe four distinct dots of CID (Centromere Identifier) corresponding to centromere pairing (Takeo *et al*. 2011; Tanneti *et al*. 2011). However, when centromeres are not all paired, we can count more than four dots. In the nuclei of *mei-W68^1/DfBSC782^*, *mei-W68^CD^* and *mei-P22^P22^*8-cell cysts, we counted an average of 4.2, 4.4 and 4.2 ± 0.9-1.1, respectively, of CID foci as compared to 4.2 ± 0.9 in the wild-type (Figure 4 A-E), indicating that most chromosomes were paired at their centromeres in these three independent mutant conditions compared to wild type germaria (two-tailed Student’s t-test, p=1 for *mei-W68^1/DfBSC782^*; p=0.7 for *mei-W68^CD^*; and p=0.8 for *mei-P22^P22^*).

**Figure 4.**
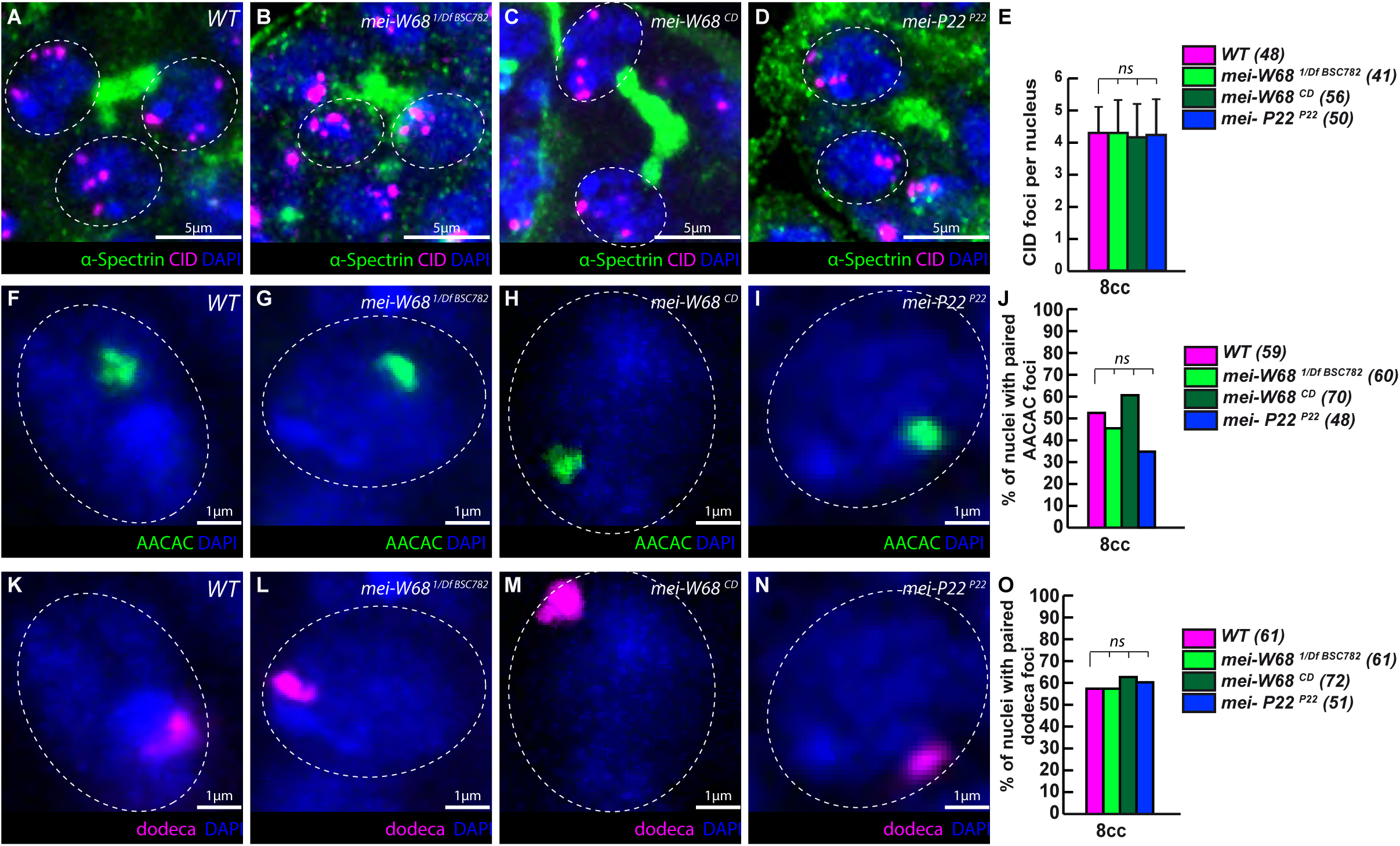
*mei-W68* and *mei-P22* are dispensable for 8-cell cyst chromosome pairing in females. A-E, Cid pairing in *mei-W68* and *mei-P22* mutant cysts. (A-D) Confocal Z-section projections of wild type (*WT*), *mei-W68^1/DfBSC782^*, *mei-W68^CD^* and *mei-P22^P22^* 8-cell cysts stained for centromeres (CID, magenta), fusome (α-Spectrin, green), and DNA (DAPI, blue). (E) Graph plots the number of CID foci per nucleus in *WT*, *mei-W68^1/Df27354^*, *mei-W68^CD^* and *mei-P22^P22^* 8-cell cysts. (*n*) is the number of cells analyzed for each genotype. ns p ≥ 0.05 (two-tailed Student’s t test comparing mutants with WT). F-O, Centromeres II and III are paired in the mitotic region of *mei-W68* and *mei-P22* mutant cysts. (F-I) Confocal Z-section projections of *WT*, *mei-W68^1/DfBSC782^*, *mei-W68^CD^*and *mei-P22^P22^* 8-cell cysts labeled with chromosome II centromeric probe (AACAC, green) and DNA (DAPI, blue). (J) Graph plots the percentage of paired chromosome II centromeres in *WT*, *mei-W68^1/DfBSC782^*, *mei-W68^CD^* and *mei-P22^P22^* 8-cell cysts. (*n*) is the number of cells analyzed for each genotype. ns p ≥ 0.05 (khi2 test comparing mutants with WT) (K-N) Confocal Z-section projections of *WT*, *mei-W68^1/DfBSC782^*, *mei-W68^CD^* and *mei-P22^P22^* 8-cell cysts labeled with chromosome III centromeric probe (dodeca, magenta) and DNA (DAPI, blue). Scale bar: 1 μm. (O) Graph plots the percentage of paired chromosome III centromeres in *WT*, *mei-W68^1/DfBSC782^*, *mei-W68^CD2^* and *mei-P22^P22^* 8-cell cysts. (*n*) is the number of cells analyzed for each genotype. ns p ≥ 0.05 (khi2 test comparing mutants with WT) Scale bar: 5 μm in A-E; 1 μm in F-O.

We also examined the pairing behavior of individual chromosomes in order to determine if pre-meiotic centromere pairing occurred between homologous chromosomes. To label the pericentromeric regions of chromosome II and III, we used the AACAC and dodeca probes respectively (Joyce *et al*. 2012). To visualize pairing, we performed fluorescence *in situ* hybridization (DNA FISH) in combination with immunostaining against the fusome marker, α-Spectrin (Figure 4F-O). We defined chromosomes as paired when only one focus was detected or when two foci were detected with a separation distance of less than or equal to 0.70 μm (GONG WJ 2005; BLUMENSTIEL JP 2008). We found that in *mei-W68^1/DfBSC782^*, *mei-W68^CD^* and *mei-P22^P22^*mutant 8-cell cysts, the number of paired chromosomes II at the level of the centromeric regions varies from 52.5% (wt) to 46.7% (*mei-W68^1/DfBSC782^*; khi^2^, p=0.5), 61.4% (*mei-W68^CD^*; khi^2^, p=0.3) and 35.4% (*mei-P22^P22^*; khi^2^, p=0.07) (Figure 4F-J); and for chromosome III, pairing varies from 57.4%, (wt) to 57.4%, (*mei-W68^1/DfBSC782^*; khi^2^ , p=1), 64.4% (*mei-W68^CD^*; khi^2^, p=0.4) and 60.8% (*mei-P22^P22^*; khi^2^, p=0.7) (Figure 4K-O). These results indicate that homologous chromosomes II and III were paired at their centromeres in all mutant conditions similarly to the wild type condition.

From these results, we concluded that Mei-W68 and Mei-P22 are not required for early centromere pairing.

### 4. Sunn, C(2)M, Nipped-B and Stromalin are dispensable for centromere pairing in the mitotic region

We previously showed that SC proteins C(3)G and Corona were expressed and required for centromere pairing in region 1 (CHRISTOPHOROU N 2013). Here, the RNAseq data indicated that many more SC or chromosome-axis proteins could be present in region 1, such as Sunn, C(2)M, Nipped-B or Stromalin (SA), which are meiotic cohesin or cohesin-associated proteins (Hughes *et al*. 2018). To test whether these genes were required for centromere pairing in region 1, we expressed shRNAs targeting each of these genes in germline cells (Figure 5). On average, we found that the numbers of centromere foci were similar between control germarium (sh-*white*) and in germarium mutant for *sunn*, *C(2)M*, *Nipped-B* and *Stromalin*, indicating that these genes are not required for the early pairing and clustering of centromeres (Figure 5A-F). We further tested the efficiency of these shRNA lines by estimating the frequencies of X-chromosome non-disjunction. We found that these lines induced efficiently between 8% and 14% of NDJ (Table S7). We concluded that, in contrast to SC proteins (C(3)G, Corona, Ord), cohesins associated to meiotic chromosomes were not required for centromere pairing.

**Figure 5.**
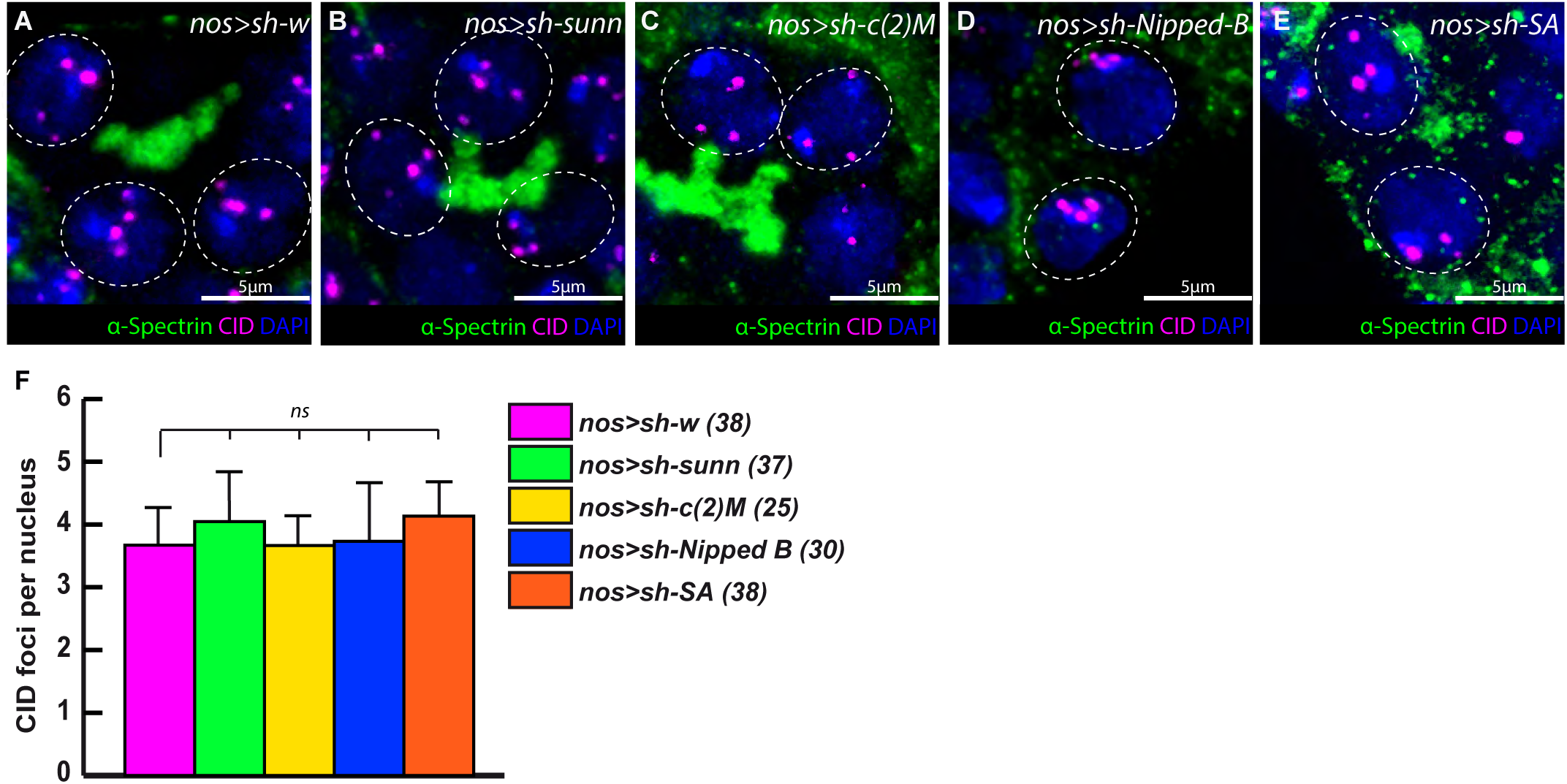
*sunn, c(2)M, Nipped-B and SA* are dispensable for 8-cell cyst chromosome pairing in females. (A-D) Confocal Z-section projections of *nos>sh-w*, *nos>sh-sunn, nos>sh-c(2)M, nos>sh-Nipped-B* and *nos>sh-SA* 8-cell cysts stained for centromeres (CID, magenta), fusome (α-Spectrin, green), and DNA (DAPI, blue). (E) Graph plots the number of CID foci per nucleus in *nos>sh-w*, *nos>sh-sunn, nos>sh-c(2)M, nos>sh-Nipped-B* and *nos>sh-SA* 8-cell cysts. (*n*) is the number of cells analyzed for each genotype. ns p ≥ 0.05 (two-tailed Student’s t test comparing *nos>sh-sunn, nos>sh-c(2)M, nos>sh-Nipped-B* and *nos>sh-SA* with *nos>sh-w* ). Scale bar: 5 μm.

### 5. DSBs activity is not detected in the premeiotic region

We then investigated whether DSBs could be present in region 1 despite the absence of Mei-W68 activity. In the *Drosophila* germline, the first sign of DSBs were described in region 2a using an antibody recognizing the phosphorylated H2A variant, also known as γH2Av (Mehrotra and Mckim 2006; Lake *et al*. 2013). To avoid using an antibody, we tested a GFP-tagged RPA transgene to label DSBs. RPA binds and protects single-strand DNA (ssDNA) just after resection of the double strand break. It is one the earliest known event of DSB repair. The coating of ssDNA by RPA is, however, transient as it is replaced by Rad51 filaments for DNA repair. To compare the pattern of DSBs precisely in the premeiotic and meiotic regions, we labeled the germarium with an antibody against α-spectrin recognizing 8cc stages, and against C(3)G to identify meiotic cells. In meiotic cysts, we selected the two pro-oocytes that displayed the brightest synaptonemal complex and counted their RPA::GFP dots in the early and late regions 2a and in region 2b (Huynh and St Johnston 2000; Page and Hawley 2001). In a wild type germarium because RPA is rapidly replaced by Rad51, the GFP signal is expected to be very rare (Figure 6). Indeed, in this genetic context (*RpA-70::GFP*), we counted an average of 0 (8 cc), 0.7 (early 2a), 2.2 (late 2a) and 0.4 (region 2b) ±0.6-1.7 GFP foci (Figure 6 A, A’, C, D, D’, G, G’, Movie S1). Furthermore, we found that most RPA-GFP foci were associated with γH2Av, while the opposite was not (Figure S7A, S7B, S7E), indicating a rapid replacement of RPA at DSBs sites.

**Figure 6.**
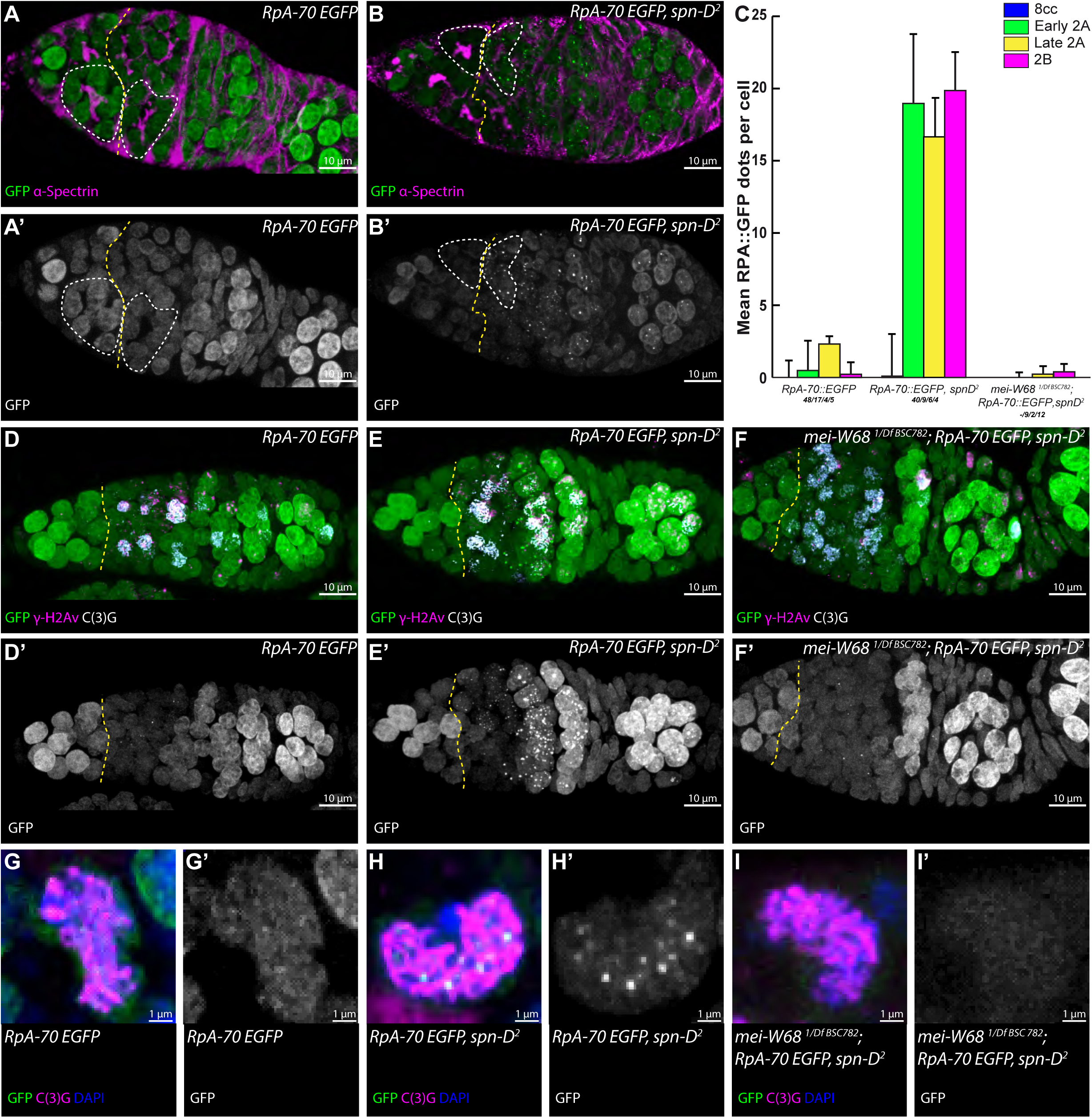
Rpa-70 foci are not detected in the pre-meiotic region of *Drosophila*. (A, A’) Confocal Z-section projection of *RpA-70 EGFP* germarium (A, green; A’, white) stained for the fusome (α-Spectrin, magenta). Note that RpA-70 EGFP is evenly distributed in the germline nucleoplasm with rare chromatin foci. Scale bar; 10 μm (B, B’) Confocal Z-section projections of *RpA-70 EGFP, spn-D^2^* germarium (B, green; B’, white) stained for the fusome (α-Spectrin, magenta). In a *spn-D^2^* mutant germarium, many RpA-70 EGFP foci are detectable in region 2 and in older egg chambers, here shown region 3. Scale bar; 10 μm. The yellow dashed line delimits the boundary of mitotic and meiotic regions, with an 8-cell cyst and a 16-cell cyst, respectively, labelled with white dashed lines. (C) Mean number of RpA-70 foci per cell counted in the mitotic 8cc (staged with α-Spectrin) and meiotic pro-oocytes (staged with C(3)G in early 2A, late 2B and 2B regions) of *RpA-70 EGFP* control*, RpA-70 EGFP, spn-D^2^* and in *mei-W68^1/DfBSC782^; RpA-70 EGFP, spn-D^2^*. The number of analyzed cells for each genetic context is labelled as *n/n/n/n*. (D-F’) Confocal Z-section projections of *RpA-70 EGFP* (D, green; D’, white), *RpA-70 EGFP; spn-D^2^* (E, green; E’, white) and *mei-W68^1/DfBSC782^; RpA-70 EGFP, spn-D^2^* (F, green; F’, white) stained for ɣ-H2Av (magenta) and for synaptonemal complex (C(3)G, white). Note that the number of RpA-70 EGFP foci in region 2 is greatly reduced in *mei-W68^1/k05603^*; *spn-D^2^* mutant cells (compare E’ and F’). Scale bar; 10 μm (G-I’) Confocal Z-section projections of selected pro-oocyte nuclei of cysts from *RpA-70 EGFP* (G, green; G’, white), *RpA-70 EGFP; spn-D^2^* (H, green; H’, white) and *mei-W68^1/DfBSC782^; RpA-70 EGFP, spn-D^2^* (I, green; I’, white) stained for synaptonemal complex (C(3)G, magenta) and DNA (DAPI, blue). Scale bar; 1 μm.

We then introduced RpA-70 EGFP into a *spn-D* mutant background (*RpA-70::GFP, spn-D^2^*). Spn-D is a meiosis-specific Rad51 homologue that is involved in removing and replacing RPA for DSBs repair in germline cells (Abdu *et al*. 2003). In this genetic context, we observed accumulation of GFP dots in the meiotic region (Figure 6B, B’, Movie S2) and not in the mitotic zone, counting an average of 0.2 (8 cc), 19 (early 2a), 16.7 (late 2a), 19.7 (2b pro-oocytes) ±0.4-4.4 GFP foci (Figure 6B, B’, C, E, E’, H, H’), respectively. We rarely detected RPA::GFP dots in the premeiotic region, indicating that neither Mei-W68 nor other sources induced detectable DSBs in the premeiotic region. In addition, we observed a much greater overlap between RPA and γH2Av dots in the *spn-D* mutant background than in the wild type condition (Figure S7C, S7D, S7E). This result confirmed the conserved role of Spn-D in RPA replacement during meiotic DSBs repair. Finally, in the additional absence of Mei-W68 (*mei-W68^1/DfBSC782^*; *RpA-70::GFP*, *spn-D^2^*), we counted 0 (8 cc), 0.2 (early 2a) and 0.3 (late 2a) ±0.3-0.4 GFP foci (Figure 6C, F, F’, I, I’), indicating that Mei-W68 is responsible for most RPA dots in a *spn-D* mutant background, and importantly, that its activity is restricted to the meiotic region.

## Discussion

In this work, we explored the transcriptome of known meiotic genes at a key transition of germline cell differentiation in *Drosophila* females. For this purpose, we used non-overlapping mitotic and meiotic cell populations genetically labelled with fluorescence transgenes in an otherwise completely wild type genetic background. Published methods for separating GSCs and differentiating cysts are based on the enrichment of GSCs in *bam* mutant conditions, and on the controlled expression of *bam* (*bam*RNAi; hs-*bam*) to enrich in differentiating cysts (Kai *et al*. 2005; Wilcockson AND ASHE 2019; Mccarthy *et al*. 2022; Samuels *et al*. 2024). Wild type ovaries have been used for single cell technology assigning differentiation stages with known markers to cell clusters (Jevitt *et al*. 2020; Slaidina *et al*. 2021). These methods have provided vast resources for functional analyses. However, they have limitations in resolving with precision the distinct stages of mitosis and meiosis: the first produces mixed population of cysts, the second generated very few cell clusters, but has expanded up to nine distinct states. Our resulting transcriptome datasets reveal that in *Drosophila*, all the genes involved in the first stages of meiosis are already expressed at low levels in the dividing germ cells before they enter the meiotic prophase I. Importantly, we were able to recover from the RNA-seq datasets known meiotic genes expressed in the mitotic compartment confirming and extending our previous findings to the whole *Drosophila* genome (CHRISTOPHOROU N 2013).

Among these genes, we confirmed by in situ hybridization that *mei-W68* is transcribed in the premeiotic region showing increasing levels in the meiotic region. These results are in agreement with previous single cell transcriptome datasets in *Drosophila* ovaries, in which germ cells in the germarium were staged using pseudotime analyses (Slaidina *et al*. 2021). Our study also provides new insights into the regulation of *mei-W68* in the germline. We inserted a small HA-His tag at the endogenous C-terminus of Mei-W68 and, although this construct is not fully functional, it allowed us to follow the pattern of *mei-W68* RNA translation. We found that Mei-W68 protein is detected mostly in early region 2a where meiotic DSBs localized and never in region 1. Thus, the primary factors contributing to the presence of Mei-W68 protein in the meiotic region are linked to the regulation of its translation. The importance of translational regulation during germ cell differentiation is well-known (Slaidina and Lehmann 2014; Teixeira and Lehmann 2019). Recently, it has been quantified genomewide using Ribo-seq, and this study showed that it is hard to predict the amount of any proteins from the corresponding mRNA levels (Samuels *et al*. 2024). Nonetheless, the presence of meiotic mRNAs in germline mitotic cells may allow a faster transition to meiosis than the activation of meiotic transcription program at the onset of meiosis.

Interestingly, in the mouse, the role of SPO11 in the initiation of pairing was recently challenged. Two independent studies found that early pairing occurred at the premeiotic stage (Boateng *et al*. 2013; Sole *et al*. 2022), while two others detected pairing at early leptotene (Ishiguro *et al*. 2014; Scherthan *et al*. 2014); however, they all agreed that early pairing events were independent of DSBs. Moreover, Boateng and colleagues further showed that pairing was dependent on SPO11 but not of its catalytic activity (Boateng *et al*. 2013). On the other hand, two independent labs found that SPO11 was not required at all for pairing (Ishiguro *et al*. 2014; Scherthan *et al*. 2014). These conflicting findings led us to ask for the requirement of Mei-W68 in premeiotic pairing in *Drosophila*. Here we show that neither Mei-W68 nor its putative partner Mei-P22 are involved in centromere pairing in the mitotic region of *Drosophila* females.

We used an Rpa70-GFP reporter as a new read-out of the initiation of meiotic recombination by DSBs (Blythe and Wieschaus 2015). Phosphorylation of the histone variant H2Av (H2AX in mammals) is a widely-used mark for DSBs (Madigan *et al*. 2002). We found that in wild-type germarium, the timing and repair of meiotic DSBs reported previously using antibodies against γ-H2Av are in agreement with our results with Rpa70-GFP. RPA foci first appeared in early region 2a, peaked in late region 2a, and then declined in 2b (Mehrotra and Mckim 2006; Lake *et al*. 2013). The number of detectable RPA foci at any one time is, however, much smaller than with γ-H2Av, confirming that RPA coating of ssDNA is very transient (Yadav and Claeys Bouuaert 2021). In contrast, in mutant conditions where DSBs are stabilized, we counted similar number of foci (19.7 in *spn-D* mutant region 2b) as previously published using antibody staining, 21.2 foci in *spn-D* mutant region 3 (Mehrotra and Mckim 2006) and 19.3 foci in *okra* mutant region 2b (Lake *et al*. 2013). As expected, in a *mei-W68* null background, no or few Rpa70-GFP foci were detected as previously reported using the γ-H2Av antibody (Mehrotra and Mckim 2006; Lake *et al*. 2013). Importantly, our results with fluorescently labeled Rpa70 confirmed that Mei-W68 does not exhibit early DSB activity in cysts before entering meiosis in region 2. Finally, our results also showed that there is no significant DSBs in the premeiotic region. In dividing embryos, the transient and rapid binding of Rpa70-GFP to sites of replication stress has allowed to measure optically the dynamics of stalled DNA replication during the mitotic cell cycle (Blythe and Wieschaus 2015). Taking advantage of the properties of this reporter, we aim to follow by live imaging the *Drosophila* germarium events of initiation and repair of DSBs in the different genetic contexts.

## Data availability statement

All fly strains are available upon request. Datasets are available from NCBI Sequence Reach Archive (SRA) under BioProject: PRJNA1011850 entitled “Isolation of stage-specific germ cells in *Drosophila* germarium”.

## Acknowledgments

We are grateful to Shelby Blythe (U. Northwestern), Bloomington Stock Center, DSHB Hybridoma Center, for antibodies and flies. We are grateful to the Orion Imaging facility at CIRB (Collège de France).

JRH lab is supported by CNRS, Inserm, Collège de France, FRM (Equipe FRM DEQ20160334884); ANR ANR-20-CE12-, BioPic) and the Bettencourt-Schueller foundation (FBS).

## MATERIAL AND METHODS

Flies were maintained on standard medium in 25°C incubators on a 12 h light/dark cycle. Wild-type controls and in combination with additional transgenes of fluorescently tagged proteins were in a *w^1118^* background.

### Fly stocks and genetics

Fly stocks used in this study: *bam::GFP/CyO; nos>UASp-RFP::wcd/TM6,tb,* is the full-length *bam* fused to GFP at C-terminus, containing its own promoter and 3’UTR (Chen and Mckearin 2003) and full-length *wicked* fused to RFP N-terminus, under the control of germline specific UASp promoter and activated by *nanos-Gal::VP16* (BDSC_4937) (Fichelson *et al*. 2009). To compare *bam::GFP/CyO; nos>UASp-RFP::wcd/TM6,tb* line to the wild type white-reference line, we first used Orb as a marker for developmental timing of germline development (Figure S1A, S1B). We found no difference between the bam-Bam::GFP; nos>wcd::RFP line and the white-control line. Orb is initially present in all germline cells in early region 1 and 2a, then becomes restricted to the oocyte in region 2b, at the anterior of the oocyte in region 3 and then switches to the posterior of the oocyte in stage 2 egg chambers. We found the two lines to be identical. We also analyzed the restriction of the synaptonemal complex to a single cell using an antibody against C(3)G (Figure S1C and S1D). We found that in region 1 and region 2a, C(3)G was identical in both genetic backgrounds. However, in region 3, we noticed that the SC signal was less intense in the future oocyte in the transgenic line (Figure S1E, “oocyte I”); and at the same time, we observed a stronger signal of C(3)G in the reverting pro-oocyte in the transgenic line (Figure S1D, open arrowhead, Figure S1E, “oocyte II”). Then at stage 2, the transgenic and control lines became identical. These data indicate that there is a transient delay in the restriction of the SC to a single cell in the *bam-Bam::GFP; nos>wcd::RFP line* compared to white-flies. We then tested whether this delay could be caused by different number of germline cysts in the germarium, but we found no difference in number of cysts in region 2 between the two genetic backgrounds (Figure S1F). We also analyzed by RNA FISH whether we could detect differences in gene expression between the two lines. We performed RNA FISH for meiotic genes found in RNAseq data, such as *c(3)G*, *Nipped-B* and *mei-W68* (Figure S3A,B,C). Quantification of FISH signals in region 2 found no difference in levels of expression of these three genes between the transgenic and control line. Finally, we used a functional assay to test for meiotic differences between these two lines, and we measured the occurrence of X-chromosome non-disjunction (Table S5). In both lines, we found only background frequencies of chromosome non-disjunctions. Overall, our thorough characterization of the *bam-Bam::GFP; nos>wcd::RFP* line revealed only a transient delay in SC restriction to the oocyte. This does not change our transcriptomic analysis of region 1 and 2.

*mei-W68 ^HA^* is a C-terminal 3x HA-linker-6x His tagged *mei-W68* fly, homozygous viable and sub-fertile generated by CRISPR/Cas9 mediated Tag knocking strategy (Well Genetics). Catalytic dead *mei-W68 ^CD^* was genome edited at the conserved catalytic domain (Y80F, Y81F) (Romanienko and Camerini-Otero 1999) using the seamless CRISPR/Cas9 strategy (Well Genetics). Flies are homozygous viable and sub-fertile. *mei-W68^1^* is a null mutation caused by spontaneous 5kb TE insertion in exon 2, females have normal synaptonemal complex but show elevated NDJ levels (Mckim and Hayashi-Hagihara 1998). Df (2R) BSC782/SM6a (BDSC_27354) is a *mei-W68* deficiency. *mei-P22^P22^* (BDSC_4931). The shRNA lines were: for *white*, P{TRiP.GL00094}attP2 (BDSC_35573); for *C(2)M* P{TRiP.GL01587}attP2 (BDSC_43977); for *SA* P{TRiP.GL00534}attP40 (BDSC_36794), for *Nipped-B* P{ TRiP.GL00574}attP40 (BDSC_36614), for *sunn* P{TRiP.HMJ21654} (BDSC_52969). *spn-D^2^* (BDSC_3326). *y w; RpA-70 EGFP[attP2]* (Blythe and Wieschaus 2015) flies were used to generate lines: *Rpa-70 EGFP spn-D^2^*, *mei-W68*^1^/CyO ; *Rpa-70 EGFP spn-D^2^*, *Df (2R)BSC782/CyO* ; *Rpa-70 EGFP spn-D^2^ /TM6,tb*.

### FACS sorted germ cells

We used the protocol for isolating mitotic and meiotic cell populations as detailed in Vallés and Huynh 2020. In brief, for each FACS isolation, 800 adult ovaries from *bam::GFP/CyO; nos>UASp-RFP::wcd/TM6,tb* flies were dissected and collected in complete medium (Schneider’s insect medium supplemented with 10% heat-inactivated fetal bovine serum, Sigma-Aldrich), dissociated with elastase at 30°C for 30 min (1 mg/ml, Sigma-Aldrich), and filtered twice (first in 40μ mesh size, then in 70μ mesh size, Corning Falcon). Cell suspensions underwent FACS separation (Aria III, BD Biosciences), collecting GFP+ and RFP+ cells and eliminating non-fluorescent cells, clumps, and dead cells. Cells were sorted directly into RNA extraction buffer (ARCTURUS Pico RNA isolation Kit, Applied Biosystems) for purification following manufacturer’s protocol.

Library preparations were done by Fasteris SA (Geneva, Switzerland) using the RNA RiboZero Stranded protocol. Indexed adapters were ligated and multiplexed sequencing performed using Illumina HiSeq 2000 (125-bp single read). At least two independent biological samples were prepared for each cell population. Sequences generated by Fasteris were aligned against the *D*. *melanogaster* reference genome (UCSC dm6) http://rohsdb.cmb.usc.edu/GBshape/cgi-bin/hgGateway.

### RT-qPCR

To validate FACS separations (Figure 1H), RFP+ and GFP+ sorted cells from *BAM::GFP; nos>Wcd::RFP* ovaries were homogenized with a pestle and RNA extracted using the ARCTURUS PicoPure RNA isolation kit.

To quantify gene expression in *mei-W68 ^1^/ Df (2R) BSC782* flies (Figure S4B), RNA was extracted from 20 pairs of dissected ovaries using the RNeasy Micro Kit (QIAGEN). RNA from *w^1118^*ovaries served as control.

For all RT-qPCR reactions, reverse transcription was done using random hexamer oligonucleotides with Superscript III Reverse Transcriptase (Invitrogen) according to manufacturer’s protocol, then by RT-PCR using *Power* SYBR Green^©^ PCR Master Mix (applied biosystems). Amplifications were done on a CFX Connect Real-Time PCR machine (Bio-Rad). Two to three biological replicates per genotype were used for all RT-qPCR experiments run in triplicate.

Relative expression levels of tested genes were calculated by the *C*t method with samples normalized to *rp49* (Schmittgen and Livak, 2008). For each experiment, primer expression in *mei-W68^1^* was compared to *w^1118^* equal 1. To compare gene expression levels between the two isolated cell populations, we first normalized each target sample (2-3) with the *C*t method (to *rp49)*. For each experiment, <u>we then normalized the highest value of the two populations to 1</u>. Expression values collected from 3-5 experiments were analyzed and transformed into graphs with Prism8 software. Mann-Whitney tests were applied to compare data.

The primers used for validation of isolated cell populations were:

GFP: F 5’AGAGGGCGAATCCAGCTCTGGAG 3’, R 5’CCCAAATCGGCGGTCAGGTGATC 3’;

RFP: F 5’ GTCCCCTCAGTTCCAGTACG 30, R 5’ TGTAGATGAACTCGCCGTC 3’;

*bam*: F 5’CTGCATATGATTGGTCTGCACGGC 3’, R 5’CCCAAATCGGCGGTCAGGTGATC 3’;

*piwi:* F 5’ CAGAGGATCTTCATCAGGTG 3’, R 5’ ATCATATTGGTCACCCCAC 3’;

*mtrm:* F 5’ GAAAGTGCCAACGAAGGTGC 3’, R 5’ CTCCATATTCGAGTCATCCGAAC 3’;

The *mei-W68* primers were:

A: F 5’ AGCTGCTGCTACTGCTGCTG 3’, R 5’ CCGACTTTTACCGAACGAAAACGAC 3’;

B: F 5’ GCTAGAACAATG GATGAATTTTCGG 3’, R 5’ GGAGAGCATGTAAATCAGCACG 3’;

C: F 5’ CGTGCTGATTTACATGCTCTCC 3’, R 5’ GACCGGACTAGCAGAGGATT3’.

*rp49*: F 5’ATCTCGCCGCAGTAAACGC 3’, R 5’CCGCTTCAAGGGACAGTATCTG 3’.

### Data analysis and heatmap generation

The DESeq2 method for differential analysis of RNA-seq data was used (Love et al 2014). As input, we used three GFP and two RFP distinct biological replicates with counts normalized for differences in sequencing depth using the DESeq normalization tool in Galaxy Mississipi^2^ platform (https://mississippi.sorbonne-universite.fr). The normalized raw counts were then used to calculate the base mean for each gene expressed in the mitotic and meiotic cell population to generate the “DESeq2 results extended with basemeans of conditions” file (Table S3). Gene lengths were taken into account by calculating FPKM for each gene (Table S4). We then extracted a subset of genes (meiotic, somatic and separation controls) and obtained Table S1 (in FPKM Table S2) used for creating a heatmap (Figure 1E). To generate the heatmap, a list of meiotic genes was compiled from FlyBase GO term (GO: 0007127), excluding genes identified as male-specific and unannotated. Added to the list are known meiotic genes (*SMC1, SMC3, sunn, solo, ord*), *RpA-70*, *dpp, egfr, Stat9e,*sorting (*bam* and *wcd*), and contamination controls possibly derived from somatic tissues like gut, fat, introduced during dissection of ovaries (*robo3, vnd, bap, twi)* (See Table S1). The resulting values were transformed to log2 and used to generate a heatmap with the heatmap2 tool in the Galaxy Mississipi^2^ platform.

### Datasets repository

Datasets are available from NCBI Sequence Reach Archive (SRA) under BioProject: PRJNA1011850 entitled “Isolation of stage-specific germ cells in *Drosophila* germarium”.

### Nondisjunction Tests

Sex chromosome nondisjunction was monitored by scoring the progeny of *y/BS Y* males mated to females carrying meiotic mutations on the second or third chromosome. For crosses with RNAi lines, the *nanos-Gal::VP16* driver was used. In most cases, a male to female ratio of 5:10 was kept. From these crosses, exceptional diplo-X and nullo-X resulting from sex chromosome nondisjunction and normal gametes are obtained. Frequency of *X* chromosome nondisjunction was calculated as 2(X-ND progeny)/total progeny, where total progeny =[2(X-ND progeny) + (regular progeny) (Gyuricza *et al*. 2016). To determine autosomal 2nd chromosome nondisjunction, females carrying meiotic mutations were mated to *C(2)EN b pr* (BDSC: 1112) males and the number of progeny scored. In most cases, a male to female ratio of 5:10 was kept. From these crosses, only the exceptional diplo-2 and nullo-2 gametes are observed.

### Immunohistochemistry

For confocal microscopy, ovaries were dissected in PBS, fixed in 4% PFA–PBS, and then permeabilized in PBT (0.2% Triton) for 30 min. Samples were incubated overnight with primary antibodies in PBT at 4 °C, washed 4 × 30 min in PBT, incubated with secondary antibody for 2 h at room temperature, washed 4 × 30 min in PBT. DAPI (1:500) was added during the last wash and then mounted in CityFluor.

For DNA FISH experiments, ovaries were dissected in PBS, fixed in 4% PFA in 1X fix buffer (100 mm potassium cacodylate, 100 mm sucrose, 40 mm sodium acetate, and 10 mm EGTA). Samples were then rinsed three times in 2X SSCT and incubated with the AACAC and dodeca probes which target the pericentromeric regions of the 2^nd^ and 3rd chromosomes, respectively, as previously described (CHRISTOPHOROU N 2013). Samples were then rinsed in 2X SSCT, twice in PBST and process for immunostaining as described above for confocal microscopy. For RNA FISH experiments, we followed the HCR *in situ* hybridization protocol for ovaries as described in (Slaidina *et al*. 2021), which was adapted from (Choi *et al*. 2018) Custom designed probes for *mei-W68* (NT_033778), hybridization buffer, wash buffer, and amplification buffer came from Molecular Instruments Inc.

The following primary antibodies were used: mouse anti-C(3)G 1A8-1G2 (1:500) (gift from S. Hawley, Stowers Institute, USA), rat anti-Cid (1:1,000) (gift from C. E. Sunkel, Universidade do Porto, Portugal), rabbit anti-α-Spectrin (1:1,000 and 1:500 when used with DNA FISH) (gift from R. Dubreuil, University of Chicago, USA), mouse anti-α-Spectrin (1:500, clone 3A9, DSHB), mouse anti-orb (1:500, clone 6H4, DSHB), mouse anti-γH2Av (1:200) (DSHB, UNC93-5.2.1), rabbit anti-HA-Tag (1:100) (Cell Signaling Technologies, C29A4).

Secondary antibodies conjugated with Cy3, Cy5, (Jackson laboratories) were used at 1:200, Alexa Fluor Plus 555, and 647 at 1:400 (Thermo Fisher Scientific).

### Image acquisition

Ovaries for imaging were taken from 3-5 day-old flies. Confocal images of fixed germaria were obtained with a Zeiss LSM 980 NLO confocal microscope except for Supplementary Figure 1. All images were acquired with a PlanApo 63×/1.4 NA oil objective at 0.5 μm intervals along the z-axis operated by ZEN 2012 software. For supplementary Figure 1, confocal images of fixed germaria were taken with a spinning-disc confocal microscope (Yokogawa) operated by Metamorph software on an inverted Nikon Eclipse Ti microscope coupled to a Coolsnap HQ2 camera (Photometrics). All images were acquired with the PlanApo 60×/1.4 NA Oil objective.

### Live imaging in oil

Ovaries were dissected in oil (10S, Voltalef, VWR) and transfer onto a coverslip. Germaria were made to stick to the coverslip in oil. All images were acquired on an inverted spinning-disc confocal microscope (Roper/Nikon) operated by Metamorph 7.7 coupled to an sCMOS camera and with a 60X/1.4 oil objective. 1 z-stack acquired every 30sec.

### Data analysis of images

For quantification of CID foci on fixed tissue, we counted the number of distinguishable CID foci within each single nucleus. In all figures, micrographs represent the projections of selected z-series taken from the first CID foci signal until the last one. For DNA FISH experiments, the 3D distances between the AACAC foci and between the dodeca foci were measured as described (CHRISTOPHOROU N 2013). Pericentromeric regions of chromosomes were considered as paired when only one foci was visible or when two foci were separated by a distance less than 0.70 μm, and as unpaired when ≥ 0.70 μm.

Fluorescence intensity measurements of RNA FISH were performed on Z-stack images acquired with identical settings. To define a Region of Interest (ROI), a Z MAX projection of 3 successive images within a circle of 50 pixels in diameter was chosen at the center of each analyzed cyst. The cyst stage was determined using the spectrin channel. Cysts located in region 2 were considered as meiotic cysts, while branched cysts of 2-cc, 4-cc, and 8-cc were classified as mitotic cysts. As background control, the ROI was selected in the somatic cells of the nascent stalk before the region 3 cyst of each analyzed germarium. For each cyst and control ROI, the Raw Integrated Density was quantified using Fiji software. The raw data were then transformed into graphs with Prism8 software. Mann-Whitney tests were used to compare fluorescence intensity.

Fluorescence intensity measurements of C(3)G were performed on Z-stack images acquired with identical settings. To define a Region of Interest (ROI), a Z MAX projection of 3 successive images within a circle of 50 pixels in diameter was chosen at the center of the C(3)G marked nuclei. The cyst stage was determined using C(3)G staining location in the germarium. As background control, the ROI was selected in the somatic cells of the nascent stalk before the region 3 cyst of each analyzed germarium. For each cyst and control ROI, the Raw Integrated Density was quantified using Fiji software. The raw data were then transformed into graphs with Prism8 software. Mann-Whitney tests were used to compare fluorescence intensity.

Mean Cyst number estimation in meiotic region 2 were performed on Z-stack images acquired with identical settings. Cysts boundaries were defined thanks to α-Spectrin staining, and counted manually.

For the quantification of RPA and H2Av foci on fixed tissues, we counted the number of distinct foci within each individual nucleus. For each channel, the signal was processed using the difference of Gaussians tool available in the GDSC plugin for Fiji. Default Threshold was then applied to the resulting stack, generating binary images reconstructed into a three-dimensional stack using the 3D segmentation function of RoiManager3D 4.1.5. The counting of RPA and H2Av dots and the percentage of ’overlap’ were then calculated using the ’Measure 3D’ analysis.

**Figure S1.**
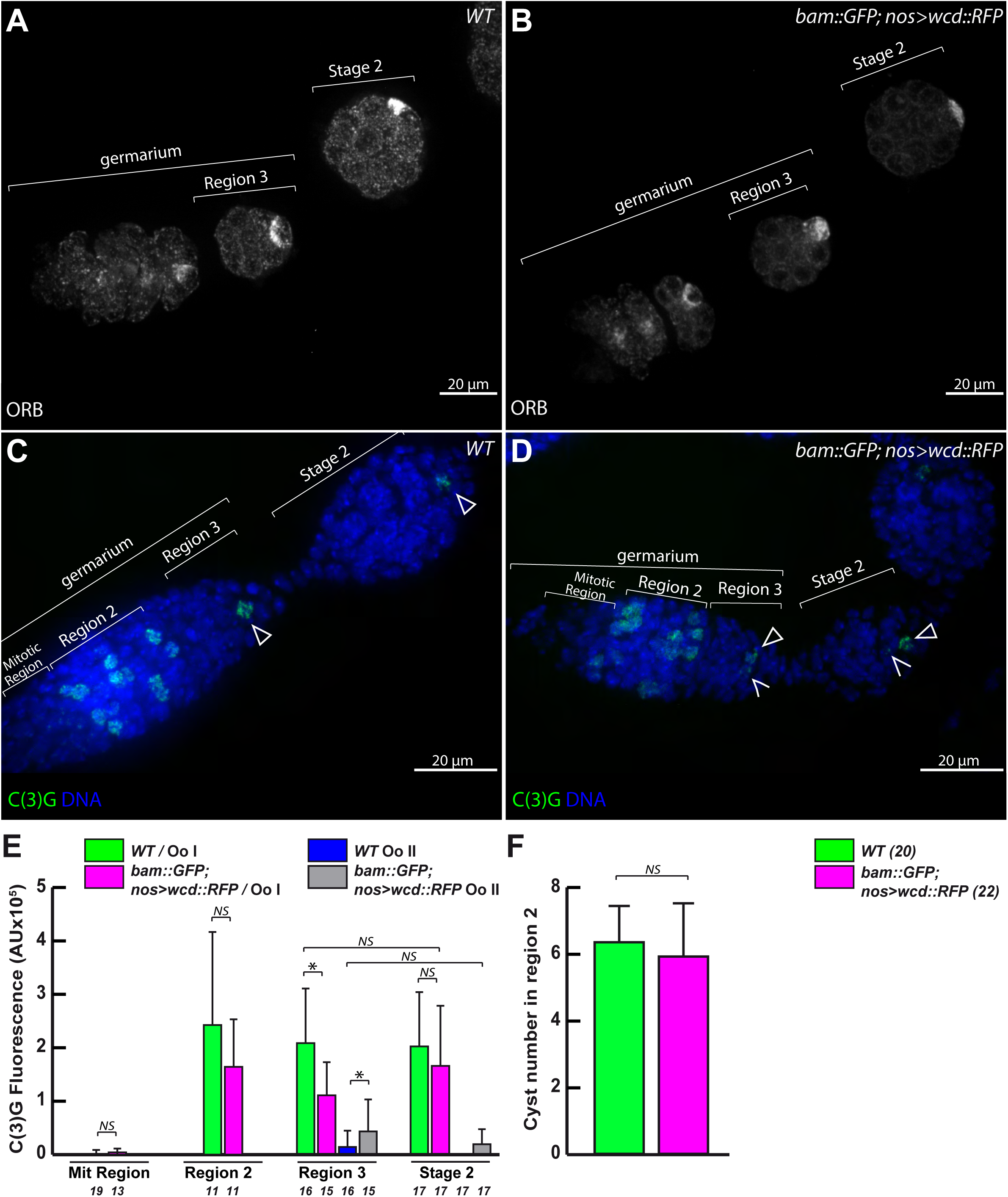
Progression of germ cells in Wcd-RFP germarium. Confocal Z-section projections of germaria from wild type (A) and bam-GFP; nos>wcd-RFP stained for ORB (white). Note that ORB localization in the meiotic region and in stage 2 is similar in both genotypes . Scale bar; 20 μm Confocal Z-section projections of germaria from wild type (C) and bam-GFP; nos>wcd-RFP (D) stained for C(3)G (green) and DNA (DAPI, blue). Note that C(3)G is restricted to the selected oocyte (oocyte I) in meiotic region 3 and stage 2 in both genotypes (arrowheads, C and D) , but the synaptonemal complex is not completely resolved in the second pro-oocyte (pro-oocyte II) in nos>wcd-RFP (open arrowheads, D). Scale bar; 20 μm (E) Graph plots C(3)G Fluorescence Intensity in the mitotic region and in the meiotic region 2 (WT, green, nos>wcd-RFP, magenta), meiotic region 3 and stage 2 (WT pro-oocyte I, green, WT pro-oocyte II , blue, nos>wcd-RFP pro-oocyte I, magenta and nos>wcd-RFP pro-oocyte II, grey). NS p ≥ 0.05, * p ≤ 0.05 (Mann-Whitney U-test). Number of samples analysed are below the bars. (F) Graph plots the number of cysts in region 2 in wild type and bam-GFP, nos>wcd-RFP calculated using spectrin antibody labelling (not shown) . NS p ≥ 0.05 (Mann-Whitney U-test). (*n*) is the number of germarium analyzed for each genotype.

**Figure S2.**
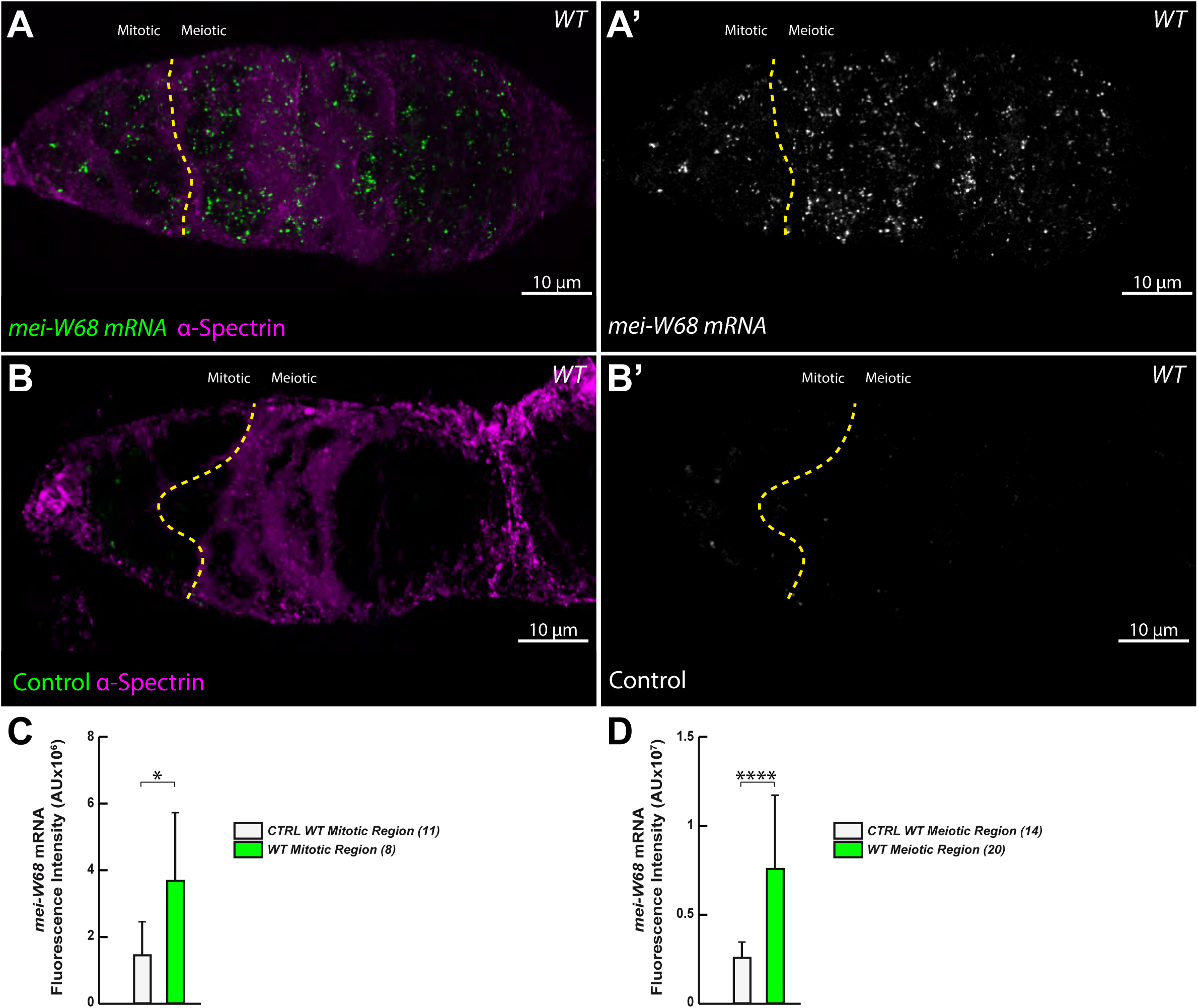
*mei-W68* mRNA signal is detected above background levels in the meiotic and mitotic regions. (A, A’) Confocal Z-section projection of a *WT* germarium labelled for *mei-W68* mRNA by HCR *in situ* hybridization using *mei-W68* HCR-initiator probes (A, green; A’, white). The yellow dashed line delimits the boundary of mitotic and meiotic regions. α-Spectrin antibody labelling is in magenta. Scale bar: 10 μm. (B, B’) Confocal Z-section projection of a *WT* germarium labelled by HCR *in situ* hybridization but excluding the HCR-initiator probes (compare with A, green; A’, white). The yellow dashed line delimits the boundary of mitotic and meiotic regions. α-Spectrin antibody labelling is in magenta. Scale bar: 10 μm. (C) Graph plots the Fluorescence Intensity in the mitotic region of WT germarium labelled by HCR *in situ* hybridization without (white) and with mei-W68 HCR-initiator probes (green). (*n*) is the number of germarium analyzed for each genotype.* p ≤ 0.05 (Mann-Whitney U-test) (D) Graph plots the Fluorescence Intensity in meiotic region of WT germarium labelled by HCR *in situ* hybridization without (white) and with mei-W68 HCR-initiator probes (green). (*n*) is the number of germarium analyzed for each genotype. **** p ≤ 0.0001 (Mann-Whitney U-test).

**Figure S3.**
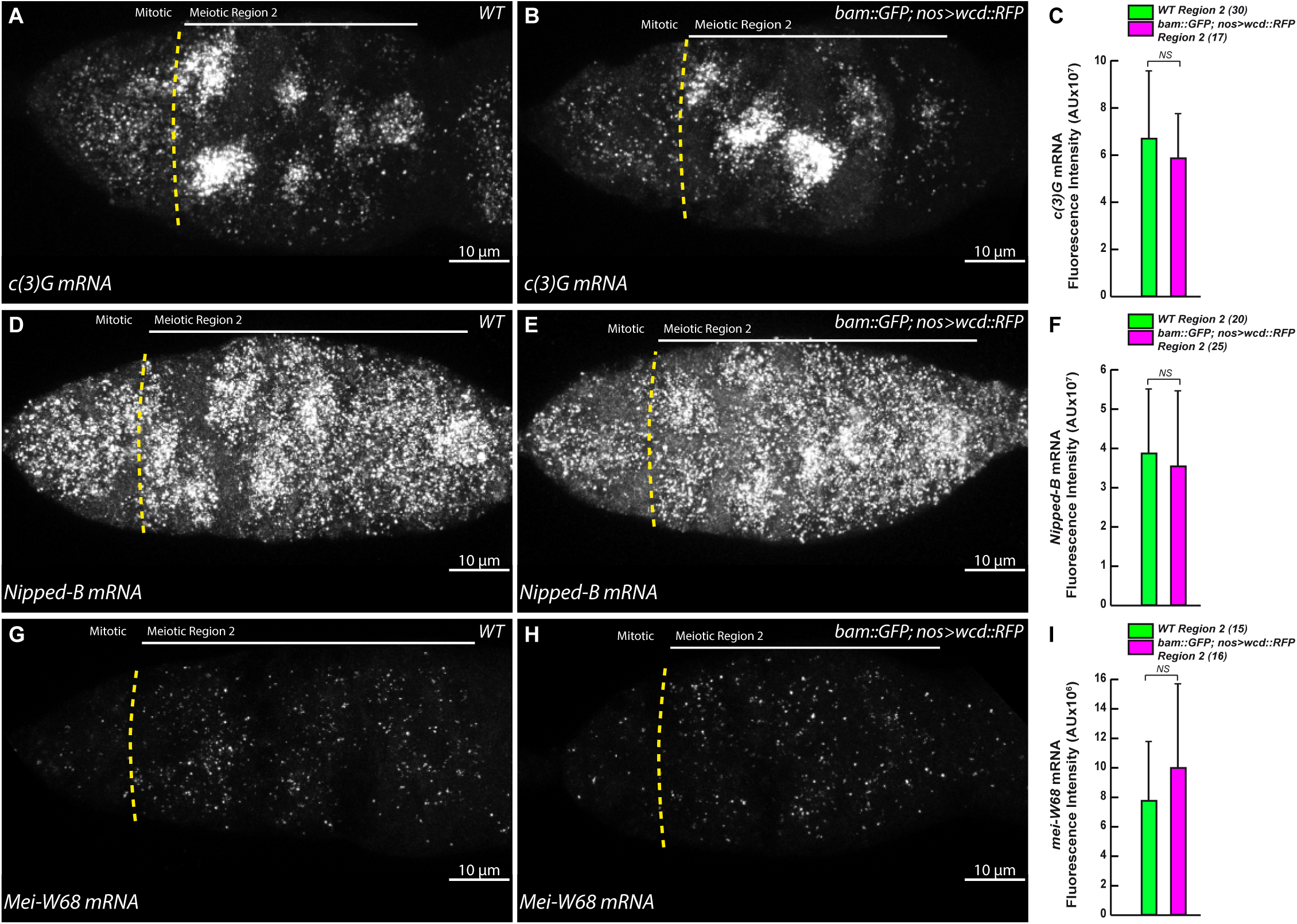
*c(3)G*, *Nipped-B* and *mei-W68* mRNA expression in the meiotic region of Wcd-RFP germarium. (A, B) Confocal Z-section projection of a *WT* (A) and *nos>wcd-RFP* (B) germarium labelled for *c(3)G* mRNA by HCR *in situ* hybridization (white). The yellow dashed line delimits the boundary of mitotic and meiotic regions. Scale bar: 10 μm. (C) Graph plots *mei-W68* mRNA Fluorescence Intensity in the meiotic region of WT (green) and *nos>wcd-RFP* (magenta) germarium labelled by HCR *in situ* hybridization. (*n*) is the number of germarium analyzed for each genotype. NS p ≥ 0.05 (Mann-Whitney U-test). (D, E) Confocal Z-section projection of a *WT* (D) and *nos>wcd-RFP* (E) germarium labelled for *Nipped-B* mRNA by HCR *in situ* hybridization (white). The yellow dashed line delimits the boundary of mitotic and meiotic regions. Scale bar: 10 μm. (F) Graph plots the *Nipped-B* mRNA Fluorescence Intensity in meiotic region of WT (green) and *nos>wcd-RFP* (magenta) germarium labelled by HCR *in situ* hybridization. (*n*) is the number of germarium analyzed for each genotype. NS p ≥ 0.05 (Mann-Whitney U-test). (G, H) Confocal Z-section projection of a *WT* (G) and *nos>wcd-RFP* (H) germarium labelled for *mei-W68* mRNA by HCR *in situ* hybridization (white). The yellow dashed line delimits the boundary of mitotic and meiotic regions. Scale bar: 10 μm. (I) Graph plots the *mei-W68* mRNA Fluorescence Intensity in the meiotic region of WT (green) and *nos>wcd-RFP* (magenta) germarium labelled by HCR *in situ* hybridization. (*n*) is the number of germarium analyzed for each genotype. NS p ≥ 0.05 (Mann-Whitney U-test).

**Figure S4.**
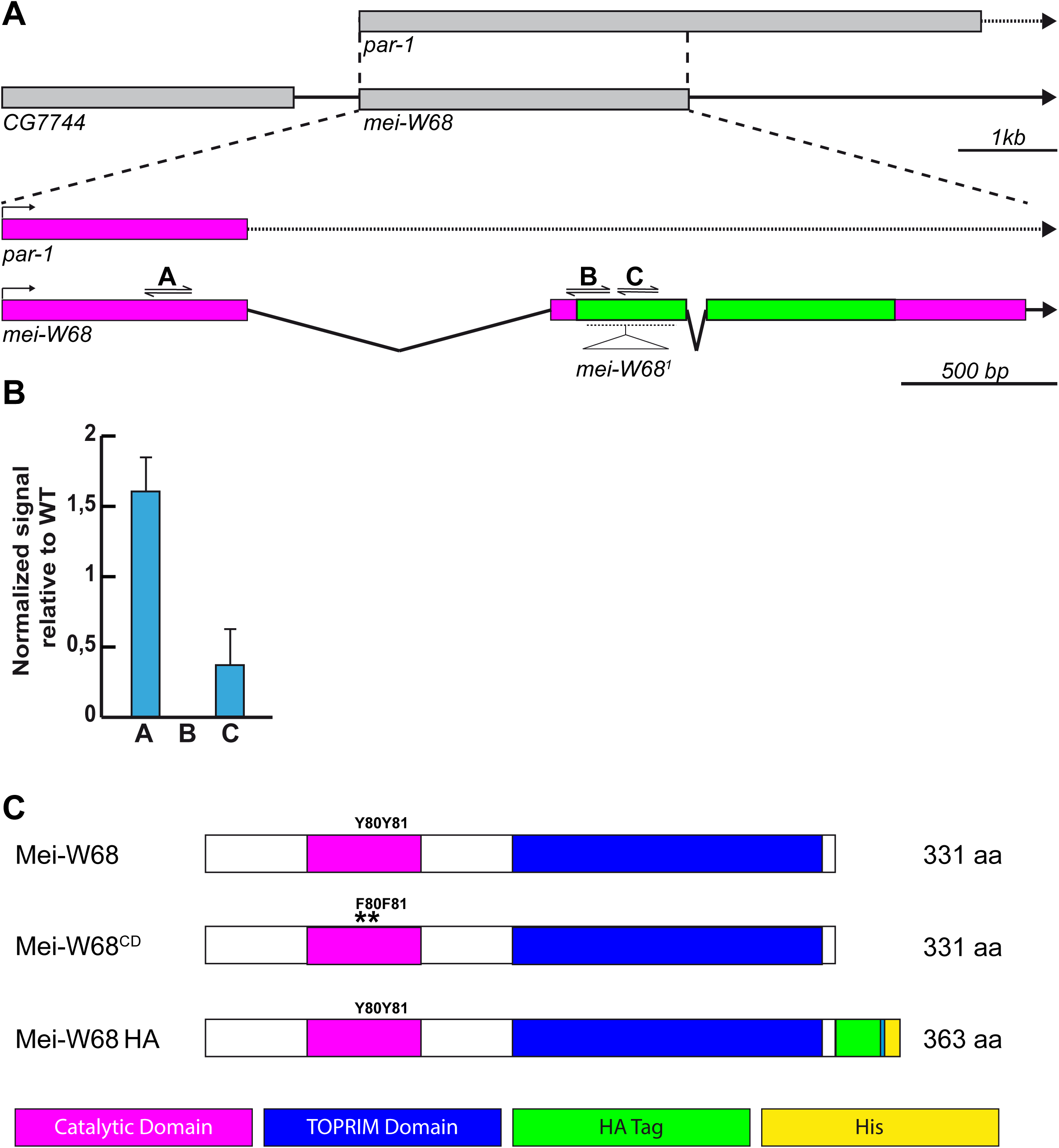
Mei-W68 reagents. (A) Schematic representation of *Drosophila melanogaster mei-W68* locus showing neighbouring genes CG7744 and *par-1* (top in grey). Enlargement of *mei-W68* and *par-1 RNA* showing introns (black acute lines for *mei-W68* and dashed lines for *par-1*), untranslated (magenta box) and translated regions (green box). Triangle represents *mei-W68^1^* insertion site of approximately 5kb (McKim and Hayashi-Hagihara). Positions of set of primers A, B and C used for RT-PCR are indicated . (B) RT-PCR gene expression levels of *mei-W68^1/DfBSC782^* using primers A, B relative to *WT*. (C) Representation of Mei-W68 catalytic (magenta box) and TOPRIM domains (blue box). Mei-W68^CD^ is a substitution in the catalytic domain of the two conserved Tyrosine (Y80Y81) into Phenylalanine (*F80*F81). Mei-W68^HA^ is tagged at the C-terminus with HA (green box) connected by a linker to His (yellow box)

**Figure S5.**
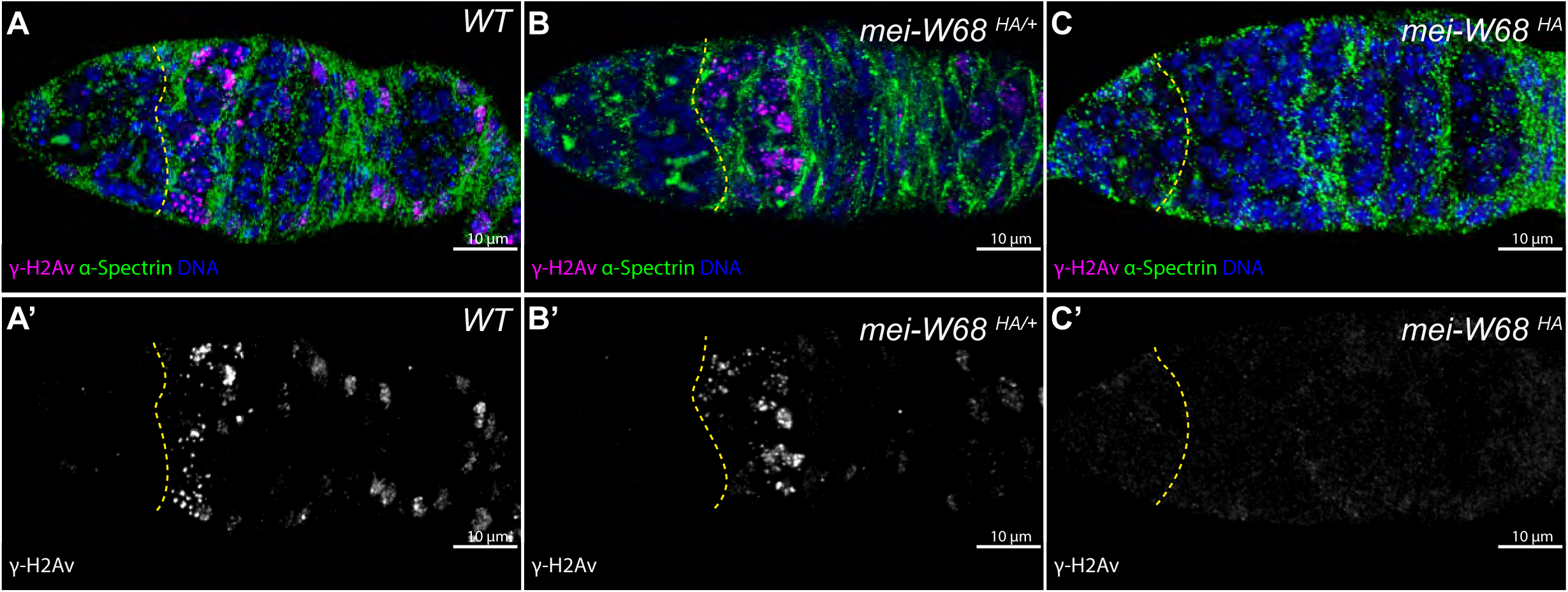
Absence of DSBs in Mei-W68^HA^ flies. Confocal Z-section projections of wild type (A, A’), *mei-W68^HA^/+* (B, B’) and *mei-W68^HA^/ mei-W68^HA^* (C, C’) germaria stained for DSBs (ɣ-H2Av, magenta), fusome (α-Spectrin, green) and DNA (DAPI, blue). The yellow dashed line delimits the boundary of mitotic and meiotic regions. Note that DSBs are absent in *mei-W68^HA^/ mei-W68^HA^*. Scale bar; 10 μm

**Figure S6.**
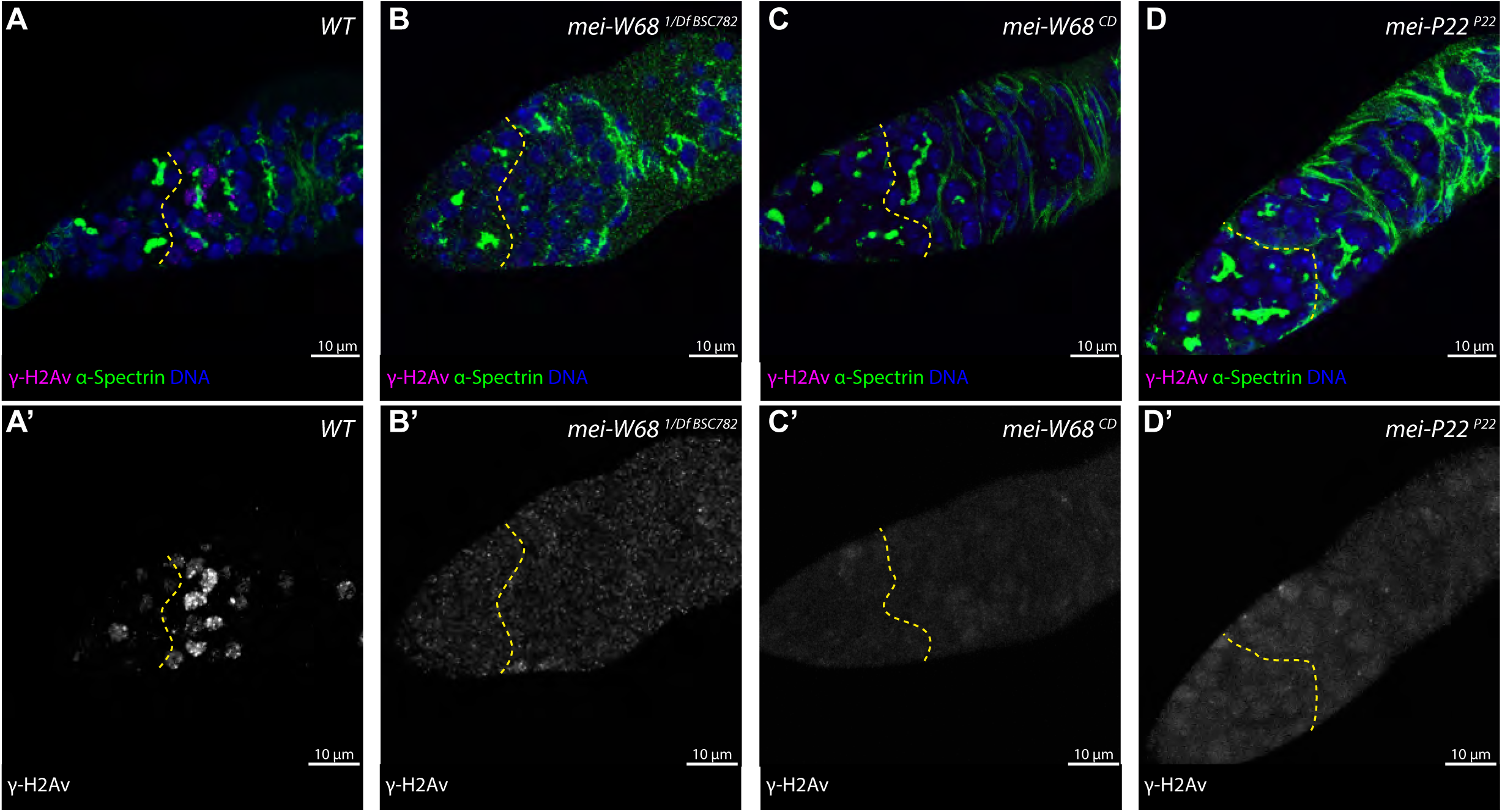
Mei-W68 and Mei-P22 mutant flies do not produce DSBs. Confocal Z-section projections of wild type (A, A’), *mei-W68^1/DfBSC782^*(B, B’), *mei-W68 ^CD/CD^*(C, C’) and *mei-P22 ^P22/P22^* (D, D’) germaria stained for DSBs (ɣ-H2Av, magenta), fusome (α-Spectrin, green) and DNA (DAPI, blue). The yellow dashed line delimits the boundary of mitotic and meiotic regions. Note that DSBs are absent in all the mutants. Scale bar; 10 μm

**Figure S7.**
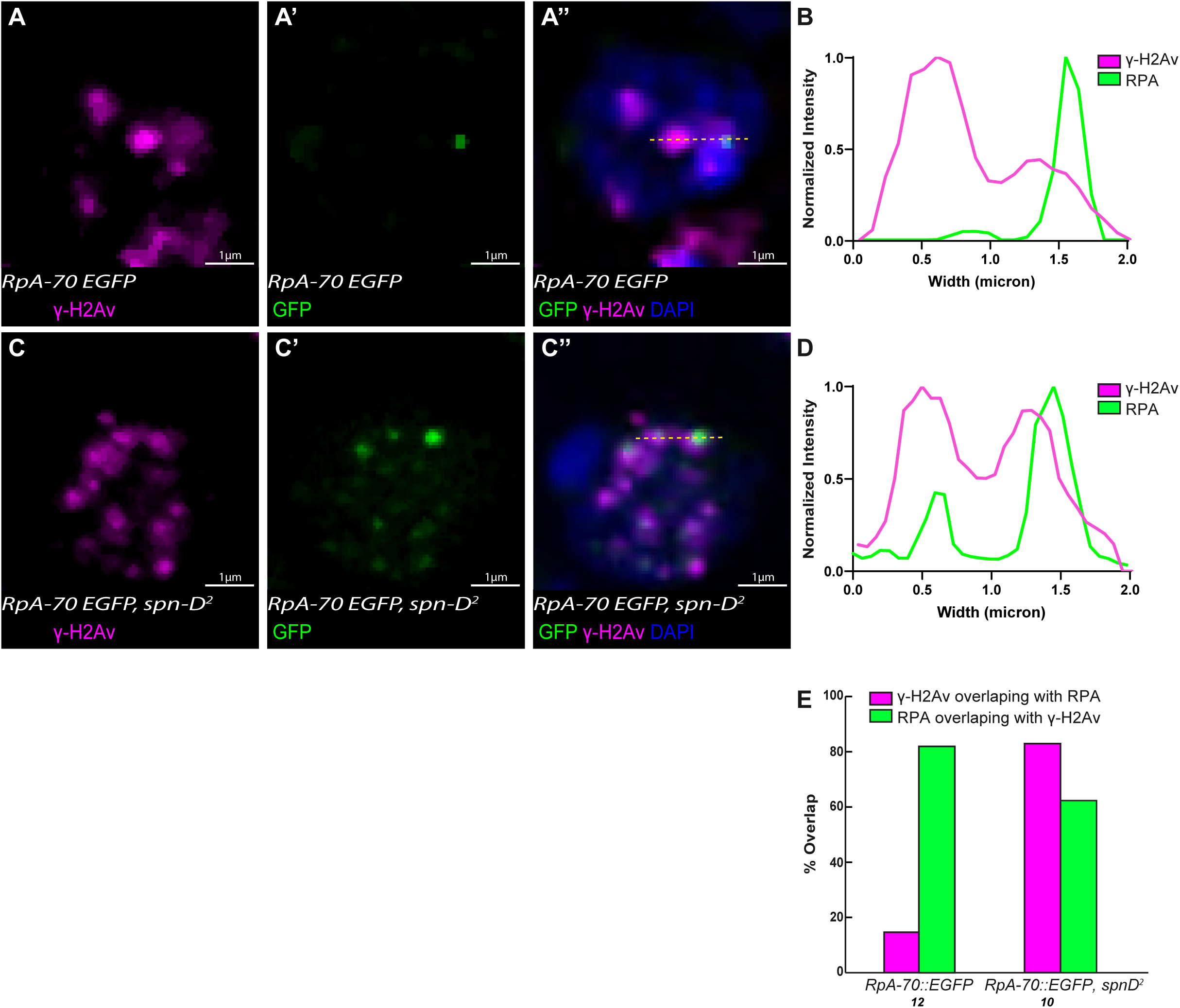
RPA::GFP foci transiently overlap with ɣ-H2Av. (A-A’’) Confocal Z-section projection of *RpA-70 EGFP* pro-oocyte stained for DSBs (ɣ-H2Av, A, A’’, magenta) and DNA (DAPI, A’’,blue). GFP is in green (A’, A’’). Scale bar= 10 μm. (B) Line profile plots the normalized intensity for ɣ-H2Av (magenta) and RPA::GFP (green) from A’’ (yellow dashed line). The RPA::GFP peak partially overlaps with the ɣ-H2Av peak on the right side, but not with the ɣ-H2Av on the left side. (C-C’’) Confocal Z-section projection of *RpA-70 EGFP; spn-D^2^* pro-oocyte stained for DSBs (ɣ-H2Av, C, C’’, magenta) and DNA (DAPI, C’’,blue). GFP is in green (C’’, C’’). Scale bar= 10 μm. (D) Line profile plots the normalized intensity for ɣ-H2Av (magenta) and RPA::GFP (green) from C’’ (yellow dashed line). The two RPA::GFP peaks overlap with the two ɣ-H2Av peaks. (E) Percentages of ɣ-H2Av overlapping with RPA::GFP (magenta) and RPA::GFP overlapping with ɣ-H2Av (green) in *RpA-70 EGFP* and *RpA-70 EGFP; spn-D^2^* pro-oocytes. The number of analyzed nuclei is indicated under each genotype.

**Movie S1**: Live-imaging of RPA::GFP in wild type germarium. Maximum intensity projection. (1 frame each 3 mn).

**Movie S2**: Live-imaging of RPA::GFP in *spn-D^2^* mutant germarium. Maximum intensity projection. (1 frame each 3 mn).

**Supplementary Table S1.**
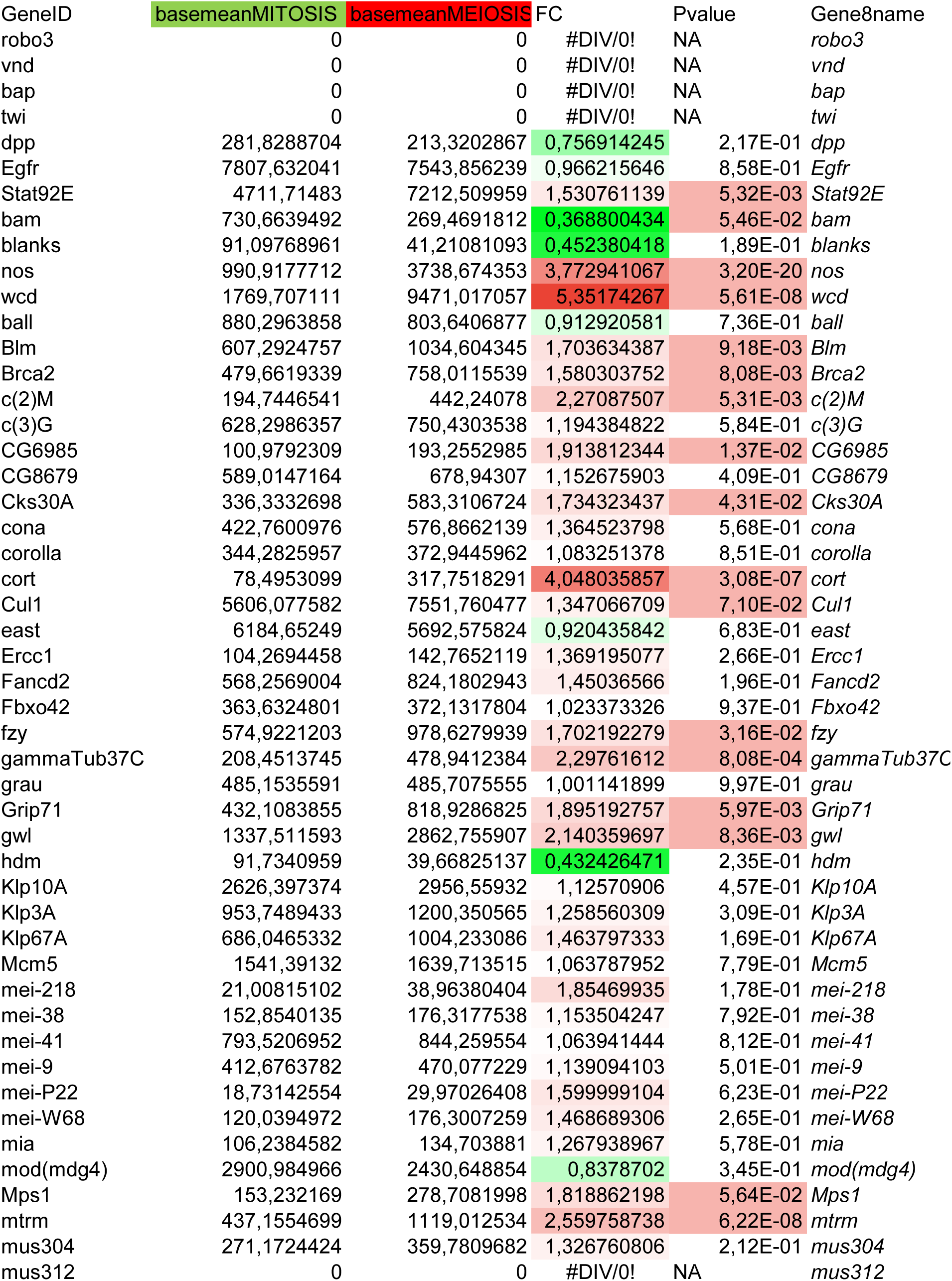
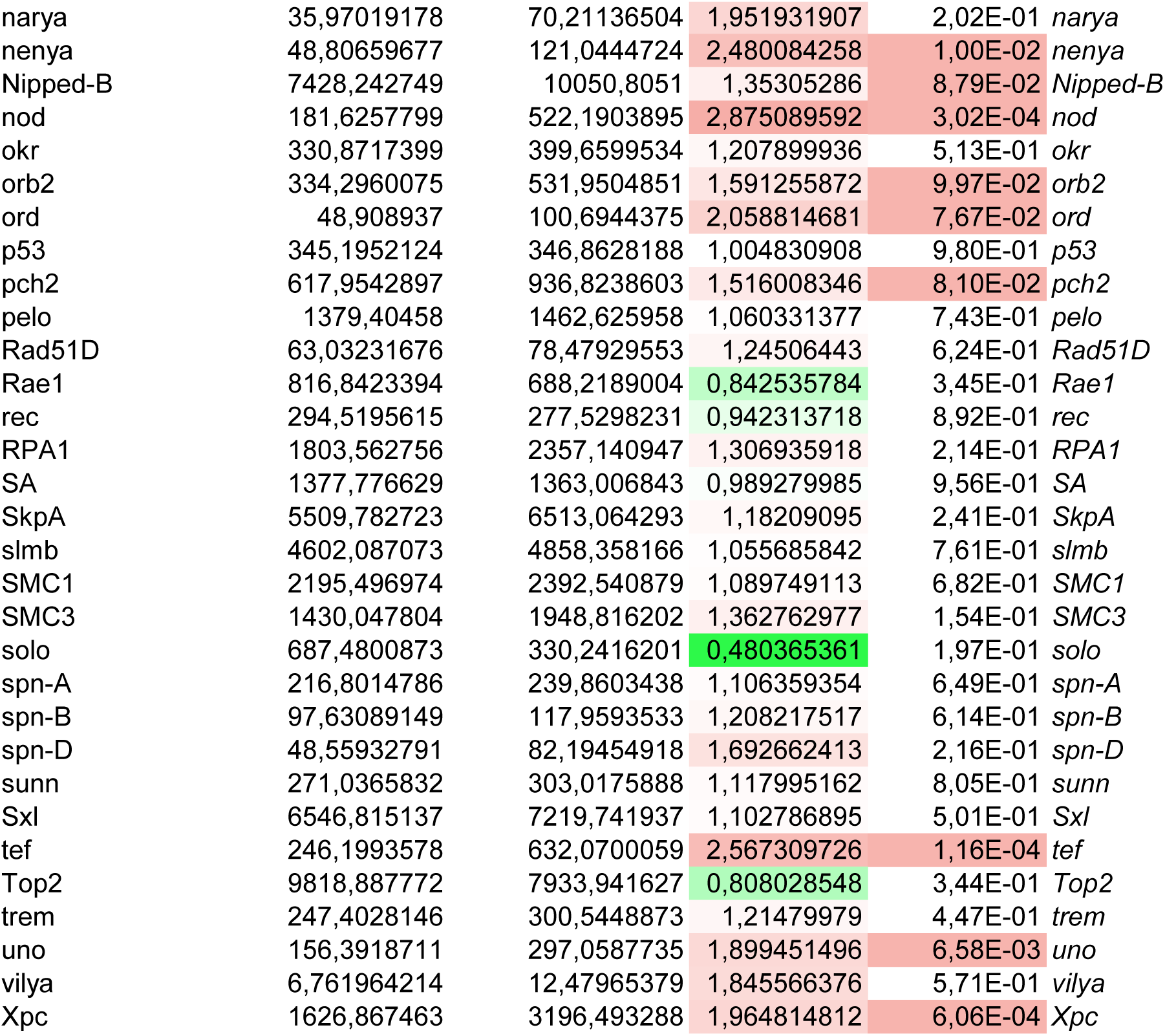
Meiotic genes are expressed in mitotic region Tab 1 - Meiotic genes compiled from FlyBase GO term (GO: 0007127), excluding genes identified as male-specific. Included in the list were sorting (bam, blanks, nos and wcd), and tissue contamination (robo3, vnd, bap, twi, dpp, Egfr and Stat92E) controls Table columns: GeneID; BaseMean_MIT; BaseMean_MEI; Fold-Change (Green <1, Red > 1); Pvalue (t-test, Red < 0,1); Gene_name

**Supplementary Table S5.** *X*-NDJ in *nos>wcd-RFP* females.

A. Sex chromosome NDJ :
| Progeny: | Regular |  | Exceptional |  |  |  |
| --- | --- | --- | --- | --- | --- | --- |
| Maternal genotype | XX | XY | XXY | X0 | Nc | % NDJ |
| *WT* | 252 | 186 | 1 | 0 | 440 | 0.5 |
| *nos>wcdRFP/+* | 212 | 204 | 0 | 1 | 418 | 0.5 |
| *nos>UAS-GFP* | 234 | 237 | 0 | 0 | 471 | 0 |

Indicated females were crossed to *w/BsYy+* males. Surviving exceptional aneuploids are Bar-eyed *y/y/B<sup>s</sup>Y* females and wild-type *yw/0* males
Percentage of X-NDJ: $100 \times 2(X\text{-NDJ progeny})/\text{total progeny}$ , where total progeny (Nc) was calculated as $2(X\text{-NDJ progeny}) + \text{regular progeny}$ (Gyuricza et al 2016).

**Supplementary Table S6.**
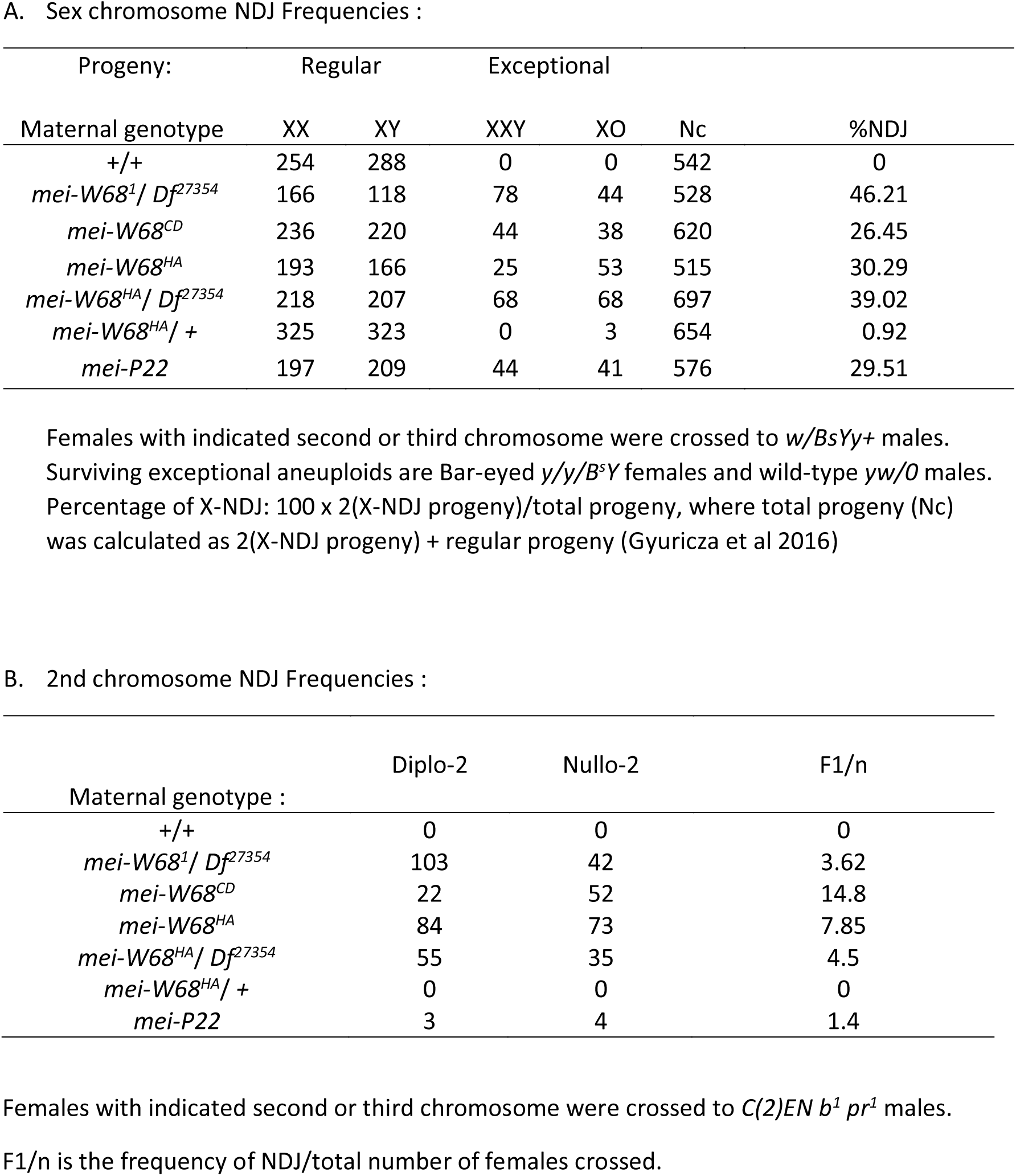
NDJ in *mei-W68* mutant females.

Sex chromosome NDJ Frequencies :
| Progeny: | Regular |  | Exceptional |  |  | %NDJ |
| --- | --- | --- | --- | --- | --- | --- |
|  | XX | XY | XXY | XO | Nc |  |
| Maternal genotype |  |  |  |  |  |  |
| +/+ | 254 | 288 | 0 | 0 | 542 | 0 |
| *mei-W68<sup>1</sup> / Df<sup>27354</sup>* | 166 | 118 | 78 | 44 | 528 | 46.21 |
| *mei-W68<sup>CD</sup>* | 236 | 220 | 44 | 38 | 620 | 26.45 |
| *mei-W68<sup>HA</sup>* | 193 | 166 | 25 | 53 | 515 | 30.29 |
| *mei-W68<sup>HA</sup> / Df<sup>27354</sup>* | 218 | 207 | 68 | 68 | 697 | 39.02 |
| *mei-W68<sup>HA</sup> / +* | 325 | 323 | 0 | 3 | 654 | 0.92 |
| *mei-P22* | 197 | 209 | 44 | 41 | 576 | 29.51 |

**B. 2nd chromosome NDJ Frequencies :**
| Maternal genotype : | Diplo-2 | Nullo-2 | F1/n |
| --- | --- | --- | --- |
| +/+ | 0 | 0 | 0 |
| *mei-W68<sup>1</sup> / Df<sup>27354</sup>* | 103 | 42 | 3.62 |
| *mei-W68<sup>CD</sup>* | 22 | 52 | 14.8 |
| *mei-W68<sup>HA</sup>* | 84 | 73 | 7.85 |
| *mei-W68<sup>HA</sup> / Df<sup>27354</sup>* | 55 | 35 | 4.5 |
| *mei-W68<sup>HA</sup> / +* | 0 | 0 | 0 |
| *mei-P22* | 3 | 4 | 1.4 |

Females with indicated second or third chromosome were crossed to *C(2)EN b<sup>1</sup> pr<sup>1</sup>* males.
F1/n is the frequency of NDJ/total number of females crossed.

**Supplementary Table S7.** *X*-NDJ in females mutant for meiotic genes.

A. Sex chromosome NDJ :
| Progeny: | Regular |  | Exceptional |  |  |  |
| --- | --- | --- | --- | --- | --- | --- |
| Maternal genotype | XX | XY | XXY | X0 | Nc | % NDJ |
| *nos>sh-w* | 393 | 169 | 0 | 1 | 564 | 0.35 |
| *nos>sh-sunn* | 322 | 313 | 29 | 24 | 741 | 14.30 |
| *nos>sh-c(2)M* | 172 | 157 | 15 | 15 | 389 | 15.42 |
| *nos>sh-Nipped-B* | 126 | 92 | 5 | 4 | 236 | 7.63 |
| *nos>sh-SA-1<sup>a</sup>* |  |  |  |  |  |  |

Indicated females were crossed to *w/BsYy+* males. Surviving exceptional aneuploids are Bar-eyed *y/y/B<sup>s</sup>Y* females and wild-type *yw/O* males
Percentage of X-NDJ: $100 \times 2(\text{X-NDJ progeny}) / \text{total progeny}$ , where total progeny (Nc) was calculated as $2(\text{X-NDJ progeny}) + \text{regular progeny}$ (Gyuricza et al. 2016)
<sup>a</sup> many embryonic lethal

